# Looping specificity of Polycomb response elements requires GAF and a combinatorial code of looping factors

**DOI:** 10.1101/2025.11.28.691166

**Authors:** Gonzalo Sabarís, Marco Di Stefano, Sandrine Denaud, Lauriane Fritsch, Ana-Maria Popmihaylova, Giorgio-Lucio Papadopoulos, Bernd Schuettengruber, Giacomo Cavalli

## Abstract

Chromatin looping between *cis*-regulatory elements is essential for precise developmental gene expression, and its disruption is frequently linked to disease. Polycomb Response Elements (PREs) are specialized tethering elements that mediate chromatin loops and are bound by transcription factors like GAGA-associated factor (GAF), contributing to the recruitment of Polycomb group (PcG) proteins. While both PcG proteins and GAF have been implicated in looping, their specific roles and the mechanisms of loop specificity remain unresolved. Using genome-wide and locus-specific approaches, we show that high GAF occupancy is required for chromatin looping and gene regulation. However, GAF alone cannot establish loops without additional factors. Surprisingly, PRE looping does not require the PcG subunit Polyhomeotic (PH) or the repressive histone marks H3K27me3. Intriguingly, orthologous PRE sequences can rescue looping, while unrelated PREs with similar GAF levels cannot. This indicates that looping specificity depends on both GAF levels and compatible factor combinations at loop anchors. Our results support a combinatorial model in which GAF collaborates with additional looping factors, to drive PRE-specific interactions. We propose the existence of a “looping code” as a mechanistic basis that might explain why only a subset of PREs form loops and contribute to Polycomb-mediated gene silencing.

## Introduction

Three-dimensional (3D) chromatin organization plays a critical role in genome function. In eukaryotes, interphase chromosomes are partitioned into physical domains (also defined as topologically associating domains or TADs), which promote local interactions between *cis*-regulatory elements (CREs) and insulate neighboring genomic regions (1–3). Recently, high-resolution Micro-C experiments in early *Drosophila* embryos identified a class of *cis*-elements named “tethering elements” (TEs) that foster chromatin loops between CREs within TADs to prime genes for rapid activation (4).

Another class of *cis*-regulatory elements forming chromatin loops and involved in the maintenance of gene expression states of key developmental genes are Polycomb Response Elements (PREs) (reviewed in (5)). PREs act as nucleation sites for the recruitment of the Polycomb repressive complexes 2 and 1 (PRC2 and PRC1), which are responsible for the deposition and spreading of repressive histone marks (H3K27me3 and H2AK118ub, respectively) forming Polycomb domains (6). These epigenetic domains often overlap with physical domains (TADs), although PcG-associated chromatin marks are dispensable for the formation of these TADs (7). Most Polycomb domains contain multiple PREs and co-regulated genes, often involved in shared developmental pathways (8,9). Only a subset of PREs forms chromatin loops (PRE loops), which are established during early *Drosophila* embryogenesis, concurrent with zygotic genome activation (10–12). PRE loops act as architectural scaffolds that form and persist independently of the underlying gene expression state (13,14). These loops can modulate gene expression by restricting enhancer-promoter communications or contributing to enhancer specificity (13). Intriguingly, a large part of the PREs correspond to TEs, with which they share common features: both elements are bound by PcG proteins, such as the PRC1 subunit Polyhomeotic (PH), and by the GAGA factor (GAF), but not by typical insulator factors such as CTCF and CP190 (4). This raises the questions of how such chromatin loops are formed and what determines their specificity, questions of increasing interest in the field.

GAF has emerged as a key player in looping formation and it is widely associated with TEs throughout the *Drosophila* genome (4,15). Similarly, it is highly enriched at PREs (12,14) and loop anchors of co-expressed genes (16). GAF possesses a BR-C, ttk and bab (BTB), or Pox virus and Zinc finger (POZ) oligomerization domain (17–19), known to mediate long-range DNA interactions (17,20). Deletion of the POZ domain results in the loss of a subset of chromatin loops, particularly between paralogous genes (15), suggesting that this domain is involved in chromatin looping between GAF bound sites. Moreover, GAF is required for the formation of a subset of the so-called, meta-loops, chromatin contacts established at the multi-megabase scale in the central nervous system (CNS) (21). Similarly, GAF can mediate long-range contacts between homologous PREs located on different chromosomes (22), and loss of GAF binding to the *dac* PRE results in reduced chromatin looping (12).

Despite this compelling evidence implicating GAF in chromatin looping, its specific function remains unclear. Only a small minority of all genomic GAF binding sites participate in loop formation, and genome-wide depletion of GAF in *Drosophila* embryos results in the loss of only a small fraction (12 out of 186) of the identified loops (15). This suggests that additional factors or features must contribute to the formation and specificity of chromatin loops. One such candidate factor is the PRC1 component PH, since looping may be mediated via oligomerization of its SAM domain. This domain is crucial for the condensation of individual Polycomb domains (23), and for mediating long-range Polycomb domain interactions in mammals (24–26). Furthermore, PREs in their transcriptionally active state (marked by lower levels of PcG proteins but retention of PH and GAF) can still form loops, suggesting a key function of these two factors for PRE looping (14).

An important experimental bottleneck in studying the direct contribution of GAF in chromatin looping is that it has a plethora of functions (reviewed in (27)) and is also implicated in PcG recruitment (12). Therefore, depleting GAF levels or analyzing GAF mutants that affect its binding affinity to chromatin, as it is the case of the POZ mutant (15), affects pleiotropic activities and prevents disentangling the direct contribution of GAF to loop formation. To overcome these limitations, we combined two complementary experimental approaches. First, we assess genome-wide correlations between GAF binding levels, motif content, and chromatin looping across development by performing CUT&RUN and Micro-C profiling in *Drosophila* embryos and larval imaginal discs. We then used CRISPR/Cas9 genome engineering to either mutate GAF binding motifs or replace a PRE in the *dac* Polycomb domain with sequence variants that differentially recruit GAF and PcG proteins, enabling us to uncouple their binding without affecting global GAF function. Our results show that, although high levels of GAF and PH are hallmarks of PRE loops, GAF is not sufficient to drive looping on its own and requires additional factors. Surprisingly, PH is also dispensable for PRE looping. Moreover, exchanging PRE sequences with homologous, orthologous or unrelated PRE sequences differentially affects PRE looping, providing the first functional evidence for a “looping code” that governs PRE loop specificity.

In analogy to PcG recruitment to PREs, which involves multiple factors, we propose that PRE looping is not governed by a single chromatin factor, but rather depends on a specific combination and dosage of multiple looping factors at loop anchors. This combinatorial code not only enables loop formation but also contributes to the specificity of looping interactions between PREs.

## Materials and Methods

### Fly work and generation of mutant flies by CRISPR/Cas9 genome engineering

All flies were raised on standard corn meal yeast extract medium at 25°C. CRISPR/Cas9 mutant fly lines ΔPRE2 and (GA)n_mut6 are described in (12). Sequences of gRNAs used to create fly lines hsp26, hsp26+3xPHO, (GA)n_mut2 and (GA)n_mut4, and PRE replacement lines (Vir, Fab-7 and en) are described in **Supplementary Table 2**. Sense and antisense oligonucleotides were annealed and phosphorylated by the T4 polynucleotide kinase (NEB#M0201S) before being inserted inside a pCFD3 plasmid (Addgene #49410) previously digested by BbsI (NEB#R0539S). pHD-dsRED donor plasmid (Addgene) containing a removable (floxed) 3XP3-dsRED construct flanked by loxP sites and DNA fragments and having homology arms to the *dac* TSS region serving as template for homology-directed repair, is described in (12). Primer sequences to insert homology arms are described in **Supplementary Table 2**.

The hsp26 sequence and the dac TSS PRE WT sequence were amplified from genomic DNA and inserted into the pHD-dsRED donor plasmid cut by SpeI and BglII using GIBSON cloning **Supplementary Table 2)**. These plasmids were used as DNA template for Multi Site-Directed Mutagenesis to generate sequences hsp26+3xPHO (STRATAGENE, #200514)) or (GA)n_mut2 and (GA)n_mut4 plasmids (Agilent Quick Change Lightning Kit #210515-5) according to manufacturer instructions. PRE sequences from the engrailed gene (en), Fab-7 or virilis PRE (vir) were amplified from genomic DNA of D. melanogaster or D. virilis and inserted into the pHD-dsRED donor plasmid cut by SpeI and BglII using GIBSON cloning **Supplementary Table 2)**.

To generate mutant fly lines, gRNA-containing pCFD3 and pHD-dsRED donor plasmids were injected into flies expressing Cas9 in the germline (vas-Cas9(X) RFP-; Bloomington stock #55821). Injections and dsRED screening was performed by BestGene (https://www.thebestgene.com/). To remove the dsRED reporter construct, mutant flies were crossed with a fly line expressing CRE recombinase (Bloomington stock #34516). Genotypes of mutant fly lines were confirmed by PCR genotyping and sequencing analysis of the mutated region.

### CUT&RUN experiments

CUT&RUN experiments were performed as described by Kami Ahmad in protocilas.io (https://dx.doi.org/10.17504/protocols.io.umfeu3n) with minor modifications. 16-20h *Drosophila* embryos were dechorionated using bleach for 2 minutes and then homogenized in 1ml of nuclear extraction buffer (20mM HEPES, 10mM KCM, 0.5mM Spermidine, 0.1% Triton X, 20% Glycerol and protease inhibitor cocktail) using a Glass douncer homogenizer. The embryo homogenates were centrifuge for 3 min at 700g and washed with nuclear extraction buffer before the addition of Concanavalin A-coated beads. 20 third instar imaginal eye or wing discs were dissected in Schneider medium, centrifuged for 3 min at 700g and washed twice with wash+ buffer before addition of Concanavalin A-coated beads. MNase digestion (pAG-MNase Enzyme from Cell Signaling) was performed for 30 min on ice. After ProteinaseK digestion, DNA was recovered using SPRIselect beads and eluted in 50ul TE. DNA libraries for sequencing were prepared using the NEBNext® Ultra™ II DNA Library Prep Kit for Illumina. Sequencing (paired-end sequencing 150bp, approx. 2Gb/sample) was performed by Novogene (https://en.novogene.com/).

### CUT&RUN Analysis

CUT&RUN data analysis was essentially performed as previously described in (28). Briefly, the sequencing read quality was assessed using FastQC and reads were aligned to the *D. melanogaster* dm6 reference genome using Bowtie 2 (v 2.4.2) (29) with the following parameters:--local---very-sensitive-local--no-unal--no-mixed--no-discordant--phred33-I 10 - X 700. The SAM alignment files were compressed into BAM files using the SAMtools (v1.16.1) software (30). Sambamba markdup (v 1.0.0) (31) was used to removed duplicate reads with parameters:-r--hash-table-size 500000--overflow-list-size 500000. The peak calling was performed with each replicate as a separate input file and IgG as the control library using MACS3 (v 3.0.0b1) (32) with the following parameters: -g dm -f BAMPE -q 0.001. Highly confident GAF peaks were defined as peaks with a MACS3 score of >100 and a fold-change of >2. Mapping and peak annotation statistics are shown in **Supplementary Table 1**. The overlap between the GAF peaks and loop anchors was determined using the GenomicRanges package in R (v 1.58.0) (33). Reads per kilobase per million mapped reads (RPKM)-normalized bigWig binary files were generated using the bamCoverage function from deepTools2 (v 3.5.5) (34). Genome browser plots were generated using the pyGenomeTracks package (v 3.8) (35) and heatmaps using the plotHeatmap function from deepTools2.

### Motif analysis

Motif scanning analysis on GAF peaks was performed using FIMO software (v 5.5.5) (36) with the MA0205.2. and GAGAGAGAGAGAGAGAGAGAGAGAGAGAG (GA29) MEME-1 GAF motif matrices. Default parameters were used and only matched sequences with a q-value ≤ 0.05 were considered.

### *k*-Means clustering of histone marks, PH profiles, and ATAC-seq data around GAF peaks

GAF peaks identified in embryo, larval eye or wing imaginal discs, or the merged dataset (for developmental comparisons; see **Supplementary Fig. 2**) were used to analyze signal tracks around the peak center. Signal intensities were normalized and stratified into a defined number of clusters using the *kmeans* function in R (version 4.4.3; www.R-project.org). Heatmaps and average signal profiles were generated using the seqplots R package (v1.23.3) (37).

### Source of CUT&RUN, ChIP-seq and ATAC-seq data

CUT&RUN data for H3K27me3, H3K27ac marks and ChIP-seq for PH data in control and PH-depleted eye imaginal discs are from (38) (GEO accession number GSE222193). The ChIP-seq data for PH in embryos was obtained from (39). The ATAC-seq data from 14-to 17-h embryos were obtained from (40), and the data for the imaginal wing discs were obtained from the 0 HS control condition in (41). The bigwig files for the two replicates were downloaded from the Gene Expression Omnibus database (accession numbers GSE120150 and GSE102839, respectively). The average of the replicates was then calculated using the bigwigAverage function from deepTools2.

### Hi-C experiments

Hi-C experiments were performed using the EpiTect Hi-C Kit (Quiagene#59971). All Hi-C experiment were performed in two or three independent experiments using 50 third instar imaginal discs. Briefly, discs were homogenized and fixed in activated Buffer T and 2% Formaldehyde using Tissue Masher tubes (Biomasher II (EOG-sterilized) 320103 Funakoshi). Tissue was digested by adding 25ul Collagenase I and II (40 mg/ml) for 1h at 37°C. Samples were centrifuged and supernatant was carefully aspirated, leaving ∼250 μl of solution in the tube. Then 250ul QIAseq Beads equilibrated to room temperature were added to bind nuclei to the beads and all subsequent reactions were performed on the beads according to the manufactures protocol. Libraries were sequenced at BGI (https://www.bgi.com/) PE 150. Sequencing statistics are summarized in **Supplementary Table 1**.

### Micro-C experiments

Micro-C experiments were performed using the Dovetail Micro-C Kit (#21006) according to manufacturer instructions with the following modifications. About 200 Drosophila embryos or 50 imaginal discs were collected or dissected into BioMasher II tube (PeloBiotech, PN: 320103) and snap frozen in liquid N2 and stored for at least 30 min at −80°C.

After thawing Embryos or imaginal discs, they were resuspended in 150ul PBS+0.3M DSG (freshly prepared) and homogenized with the pestel provided with the BioMasher II tubes. After material is completely homogenized, additional 850ul PBS+0.3M DSG is added and tubes are rotated at room temperature for a total time of 10 min. Then 27ul of 37% Formaldehyde are added for additional 10 min. After fixation, nuclear pellet is washed twice with 1ml 1x Wash Buffer. After the second wash, the pellet is resuspended again in 1ml 1x wash Buffer and using a 1 mL syringe, gently pushed through a 50μm filter into a new 1.5 mL tube. After centrifugation, the nuclear pellet is resuspendend in 50ul 1X Nuclease Digest Buffer. Embryonic nuclei were digested for 15min at 22°C with 0.5ul MNAse (undiluted), whereas nuclear pellet from imaginal discs was digested with 0.5ul MNAse (1:10 diluted). Mnase digestion was stopped by adding 5ul 0.5M EGTA. For embryo Micro-C about 1500ng of the lysate was used as Input for Micro-C procedure. For disc Micro-C the whole lysate was used.

Library preparation was performed using theNEBNext® Ultra™ II DNA Library Prep Kit for Illumina (12 cycles, DNA Polymease from Dovetail Kit). All Micro-C experiment were performed in duplicates. Libraries were sequenced at BGI (https://www.bgi.com/) PE 100. Sequencing statistics are summarized in **Supplementary Table 1**.

### Hi-C and Micro-C analysis

Raw data from Hi-C sequencing were processed using the ‘scHiC2’ pipeline and from Micro-C with a modified version to filter-out contacts at distance separation lower than 200 bp. The samples for Embryo_WT, WD_WT, (GA)n_mut and larvae_DWT were aligned to the *D. melanogaster* reference genome dm6 in https://s3.amazonaws.com/igenomes.illumina.com/ and samples for the *dac* PRE2 mutants (DPRE2, Vir, 2xPRE1, Fab-7, and en) were aligned to a modified genome. Briefly, the 644 bp region chr2L:16,485,929-16,486,572 including the PRE2 was removed from the dm6 reference genome and replaced with strings corresponding to the specific mutation (**Supplementary Table 2**). For all mutations except for 2xPRE1, we added a sequence of N (indetermined) nucleotides to replace entirely the removed sequence of 644. For 2xPRE1 mutant, we cut the end of chr2L by 459 bp to maintain its total length of 23,513,712 bp, as for the WT. To improve mappability, the *.fastqs* of Micro-C reads were trimmed to 100nt (if necessary) using trim_galore (https://github.com/FelixKrueger/TrimGalore). Sequencing statistics are summarized in **Supplementary Table 1**. Valid-pairs were stored in a database using the ‘misha’ R package (https://github.com/msauria/misha-package). Extracting the valid interactions from the misha database, the ‘shaman’ R package (https://bitbucket.org/tanaylab/shaman) was used for computing the Hi-C expected models and Hi-C scores with parameters k = 250 and k_exp = 500 (**Figs. 2c, 2e**, **3b**, **4e**, **5b**, **5d**, and **Supplementary Fig. 4b**). Specifically, Hi-C scores quantify the contact enrichment (positive values) or depletion (negative values) of each bin of the map with respect to a statistical model used to evaluate the expected number of counts. To generate this expected model, we randomized the observed Hi-C contacts using a Markov chain Monte Carlo-like approach per chromosome (42)). Shuffling was conducted such that the marginal coverage and decay of the number of observed contacts with the genomic distance were preserved but any features of genome organization (for example, TADs or loops) were not. These expected maps were generated for each biological replicate separately and contained twice the number of observed cis contacts. Next, the score for each contact in the observed contact matrix was calculated using the k nearest neighbors (kNN) strategy (42). In brief, the distributions of two-dimensional Euclidean distances between the observed contact and its nearest k_exp neighbors in the pooled observed and pooled expected (per cell type) data were compared, using Kolmogorov–Smirnov D statistics to visualize positive (higher density in observed data) and negative (lower density in observed data) enrichments. These D scores were then used for visualization (using a scale from −100 to +100) and are referred to as Hi-C scores in the text. Accordingly, the color scale of the Hi-C scores comprises both positive and negative values. For each condition, the Hi-C and Micro-C interaction quantifications at PRE loops (**Figs. 2d**, **2f**, **3c**, **4f**, **5c**, **5e**, **Supplementary Fig. 4c**, and **8c**) were performed by considering the Hi-C scores between two regions of 6 kb, chr2L:16,419,514–16,425,515 and chr2L:16,482,929–16,488,930), including the *dac* PRE1 and PRE2 or two regions of 6 kb, chrX:14,650,251-14,659,252 and chrX:14,748,251-14,754,251 including the NetA/NetB PREs (**Fig. 2e**), respectively. To measure the total enrichment of contacts, between pairs of GAF peaks whose centers are separated by at least 10 kb and not more than 500 kb (**Fig 1c**, **Supplementary Fig 2b**, and **3a-b**) and identified loops an enrichment analysis was applied. Briefly, the log_2_(Observed/Expected) score in squared windows of size *w* centered at the considered features (*w*=20 kb for GAF peaks and *w*=8 kb for Micro-C loops) at 250 bp resolutions was computed using the *shaman_generate_feature_grid_2d* function of the Shaman package. The box plots in the distributions of Hi-C and PE-scan enrichment scores (**Fig 1c,d**, **3d, Supplementary Fig**. **2b**, and **3a-b**) show the median (central line), the 75th and 25th percentiles (box limits) and 1.5 × the interquartile range (IQR; whiskers). Statistical comparisons between distributions of Hi-C and contact enrichment scores per conditions were performed using the Kruskal–Wallis test followed by Dunn’s post-hoc pairwise test with Benjamini–Hochberg correction. *Adjusted p-values* < 0.05 are reported on top of the compared distributions.

**Fig. 1:**
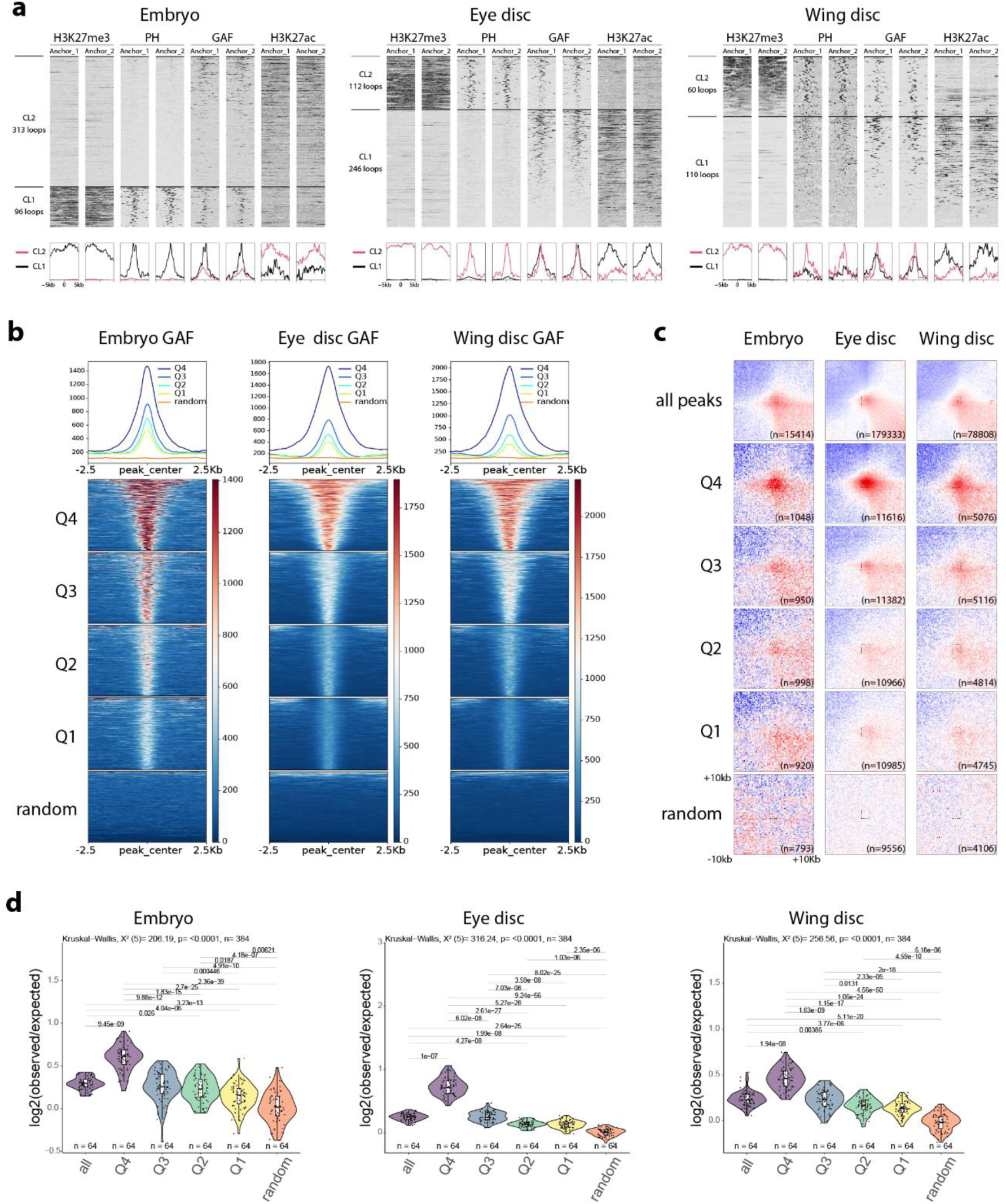
GAF binding strength correlates with chromatin looping. **(a)** Heatmaps and average signal profiles showing *k*-means clustering of H3K27me3, PH, GAF and H3K27ac signals around the anchors (1 and 2) of the loops detected in Micro-C of embryos (left) larval eye discs (middle) and larval wing discs (right). **(b)** Heatmaps and average signal profiles of GAF CUT&RUN (two merged replicates) centered on GAF peaks ranked and grouped into quartiles based on the MACS3 peak score. Random peaks matched in number and average size were used as control. **(c)** Aggregate Micro-C map at 250 bp resolutions around pairs of GAF peaks, which are grouped as in panel (b) at distance separation between 10 and 500 kb and quantification of the signal in the central 8×8 square. **(d)** Boxplots show median (central line), Q1=25th and Q3=75th percentiles (box limits), and Q1+1.5×IQR to Q3+1.5×IQR (whiskers), where IQR is the interquartile range. Statistical comparisons were performed using the Kruskal–Wallis test followed by Dunn’s post-hoc pairwise test with Benjamini–Hochberg correction.

**Fig. 2:**
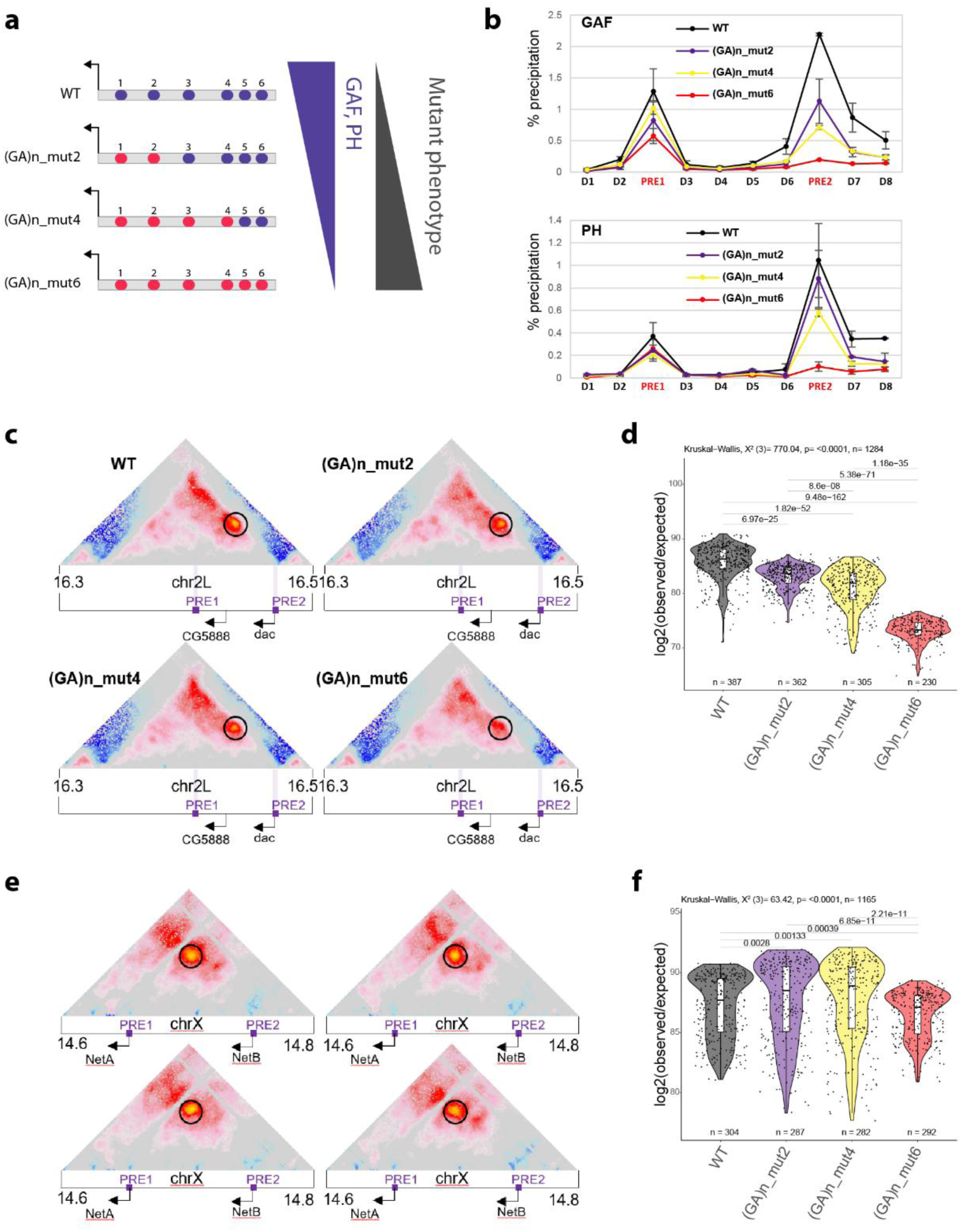
GAF binding levels are a major determinant for PRE loop formation. **(a)** Schematic representation of the PRE2 with (GA)n motifs (GAF binding sites) shown in blue. Mutated motifs are indicated in red *(left)*. Schematic representation of the consequences of mutating increasing numbers of (GA)n motifs on the binding of indicated chromatin factors (*right*). **(b)** qChIP experiments using GAF or PH antibodies in WT, (GA)n_mut2, (GA)n_mut4 and (GA)n_mut6 fly lines. D1-D8, PRE1, 2 indicate PCR amplicons used for qChIP experiments along the *dac* gene locus (Supplementary **Table 2**). The housekeeping gene *Rp49* was used as negative control, the *engrailed* PRE as a positive control. Data are presented as the mean values ± s.d (error bars) of two independent replicates. **(c, e)** Micro-C score (**Material and Methods**) maps of a 200 kb region at 1kb resolution on chromosome 2L at *dac* gene locus **(c)** and on chromosome X at the *NetA/B* gene locus **(e)** in WT, (GA)n_mut2, (GA)n_mut4 and (GA)n_mut6 fly lines. Black circles indicate the positions of the PRE loop. Violet bars indicate position of PREs. Black arrows indicate gene promoters of the *dac*, *CG5888*, *NetA*, and *NetB* genes. **(d, f)** Quantification of the Micro-C *dac* **(d)** and *NetA/B* **(f)** PRE loop interaction scores in WT, (GA)n_mut2, (GA)n_mut4 and (GA)n_mut6 fly lines. Violin plots show median (central line), Q1=25th and Q3=75th percentiles (box limits), and Q1+1.5×IQR to Q3+1.5×IQR (whiskers), where IQR is the interquartile range. Statistical comparisons were performed using the Kruskal–Wallis test followed by Dunn’s post-hoc pairwise test with Benjamini–Hochberg correction.

**Fig. 3:**
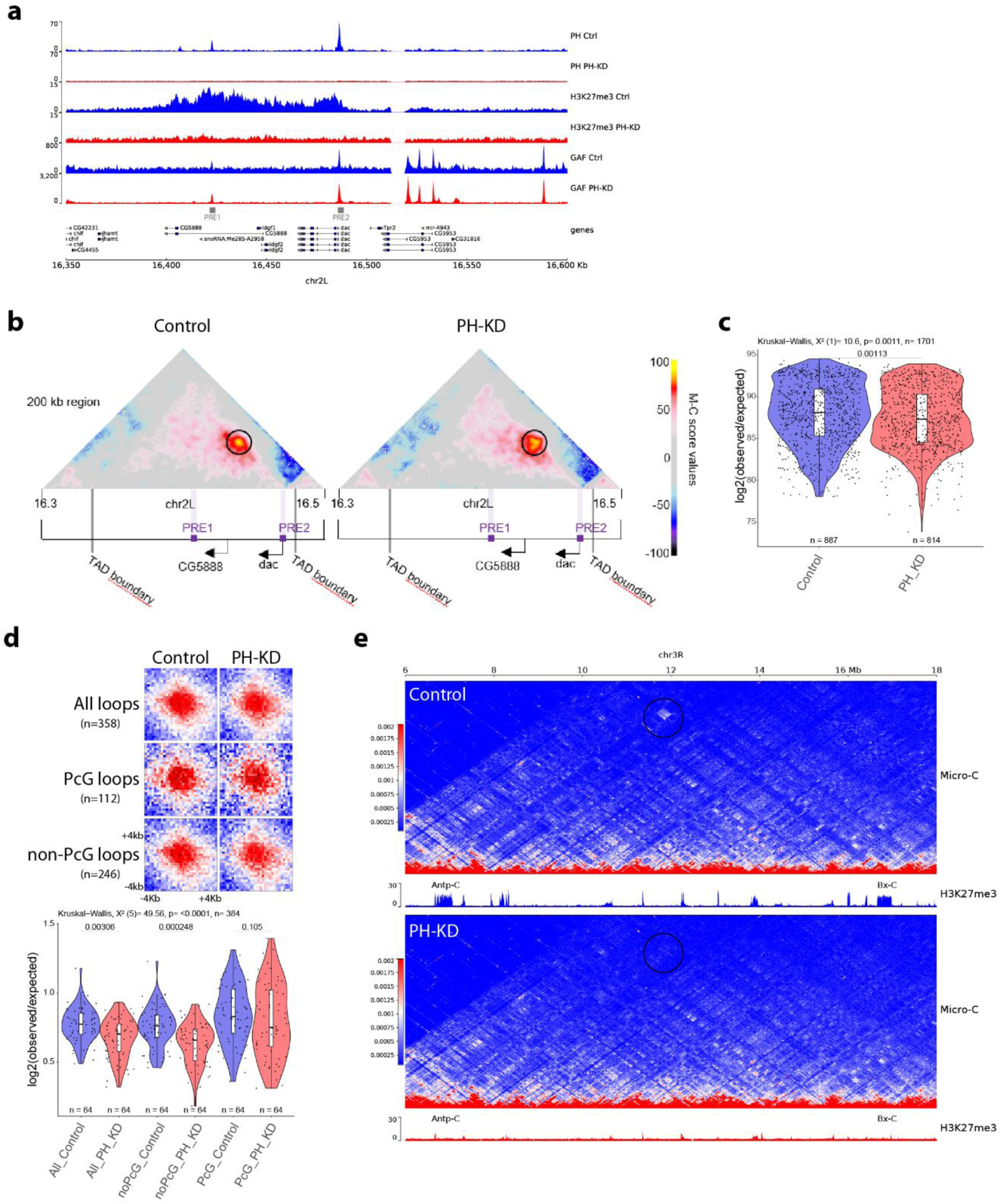
PH is not necessary for PRE looping. **(a)** ChIP-seq profiles for PH and CUT&RUN profiles H3K27me3 and GAF in Control (Ctrl) or PH-depleted (PH-KD) larval eye imaginal discs at the *dac* gene locus. **(b)** Micro-C score maps of a 200kb region around the *dac* TAD in control or PH depleted (PH KD) larval eye imaginal discs. PRE loop is indicated by a black circle. Violet bars indicate position of PREs. Gray bars indicate the positions of TAD boundaries. Black arrows indicate the promoters of the *dac* and the *CG5888* genes. **(c)** Quantification of the *dac* PRE loop Micro-C interaction scores in Control and PH depleted (PH-KD) mutant flies. Statistical comparisons were performed using the Kruskal–Wallis test followed by Dunn’s post-hoc pairwise test with Benjamini–Hochberg correction. **(d)** Aggregate Micro-C map at 250 bp resolutions centered at *All*, *PcG*, and *non-PcG* loops in Control and PH depleted (PH-KD) Micro-C (see **Supplementary Table 1** and **Material and Methods**). Violin plot shows the quantification of the signal in the central 8×8 square of the aggregate Micro-C maps. Statistical comparisons were performed using the Kruskal–Wallis test followed by Dunn’s post-hoc pairwise test with Benjamini–Hochberg correction. Boxplots in panels **(c, d)** show median (central line), Q1=25th and Q3=75th percentiles (box limits), and Q1+1.5×IQR to Q3+1.5×IQR (whiskers), where IQR is the interquartile range. **(e)** Micro-C contact maps in control eye discs (top) or upon loss of PH function (PH-KD, bottom) showing the loss of long range contacts between the Antp-C and the Bx-C (black circle).

**Fig. 4:**
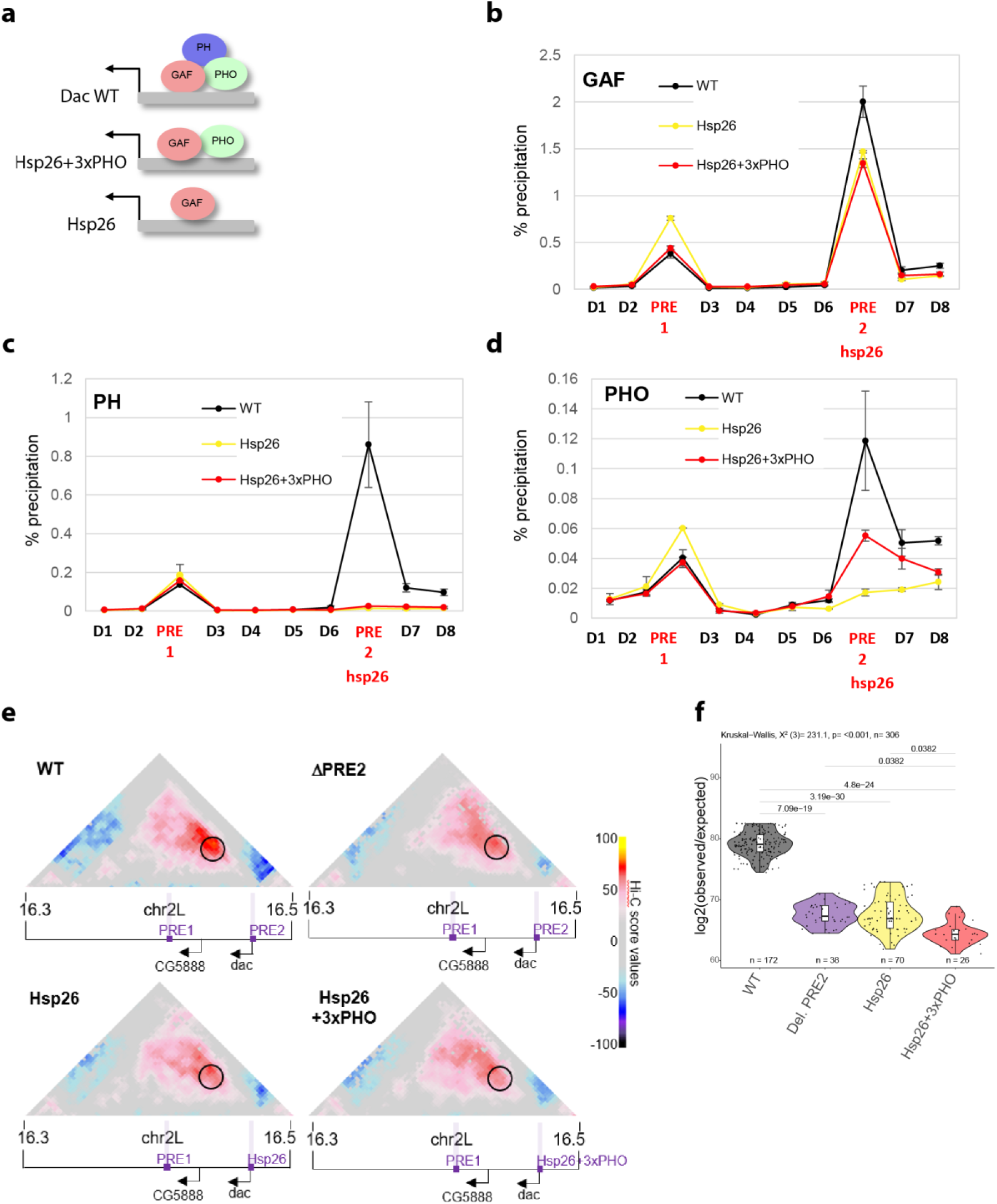
GAF is not sufficient for PRE loop formation. (**a**) Schematic representation of the Hsp26 constructs inserted at the *dac* TSS (PRE2) genomic region. **(b-d)** qChIP experiments using GAF, PH or PHO antibodies in WT, Hsp26, and Hsp26+3xPHO fly lines. D1-D8, PRE1, 2 indicate PCR amplicons used for qChIP experiments along the *dac* gene locus (Supplementary **Table 2**). Note that, to discriminate endogenous *hsp26* sequence from the ectopically inserted sequence, we used a primer pair, where one primer is specific to the *hsp26* sequence whereas the other one is specific to the *dac* promoter region. The housekeeping gene *Rp49* was used as negative control, the *engrailed* PRE as a positive control. Data are presented as the mean values ± s.d (error bars) of two independent replicates. **(e)** Hi-C score (**Material and Methods**) maps of a 200 kb region at 3kb resolution on chromosome 2L at *dac* gene locus in imaginal disc in WT, PRE2 deleted (ΔPRE2), Hsp26, and Hsp26+3xPHO fly lines. Black circle indicates the position of the *dac* PRE loop. Violet bars indicate position of PREs. Black arrows indicate gene promoters of the *dac* and the *CG5888* genes. **(f)** Quantification of the *dac* PRE loop interaction scores. Hi-C interaction score in WT, PRE2 deleted (ΔPRE2), Hsp26, and Hsp26+3xPHO fly lines. Boxplots show median (central line), Q1=25th and Q3=75th percentiles (box limits), and Q1+1.5×IQR to Q3+1.5×IQR (whiskers), where IQR is the interquartile range. Statistical comparisons were performed using the Kruskal–Wallis test followed by Dunn’s post-hoc pairwise test with Benjamini–Hochberg correction.

**Fig. 5:**
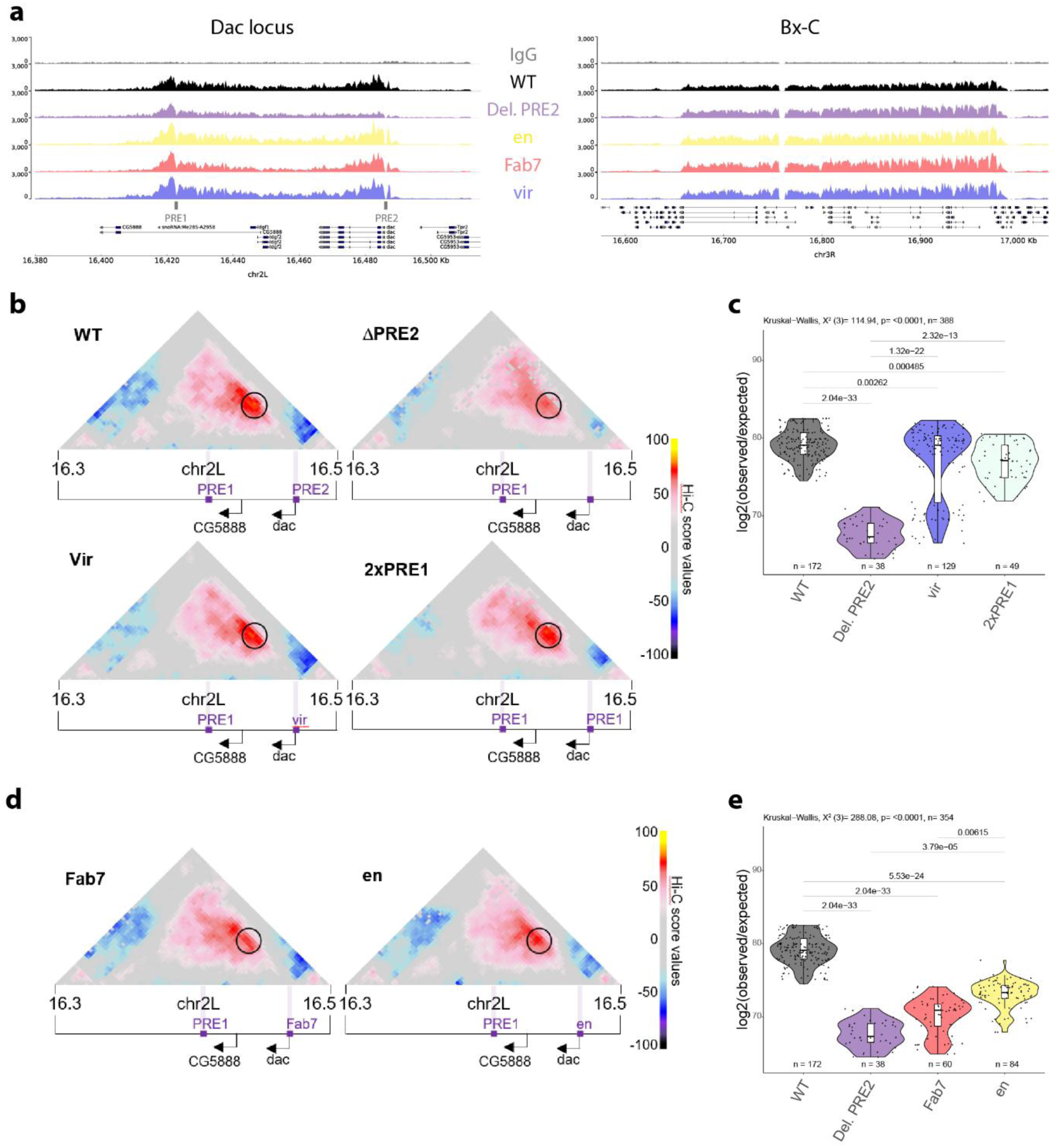
An orthologous PRE sequence can rescue loss of PRE looping. **(a)** CUT&RUN profiles of H3K27me3 around the *dac* gene (left) or the Bx-C locus (right) in WT, PRE2 deleted (ΔPRE2), or mutant PRE lines, where the *dac* PRE2 has been replaced by the indicated PREs (en, Fab-7 or vir). **(b, d)** Hi-C score (**Material and Methods**) maps of a 200 kb region at 3kb resolution on chromosome 2L at *dac* gene locus in imaginal disc in WT, PRE2 deleted (ΔPRE2), or mutant PRE lines, where the *dac* PRE2 has been replaced by the indicated PREs (vir, 2xPRE1 and Fab-7, en). Black circle indicates the position of the *dac* PRE loop. Violet bars indicate position of PREs. Black arrows indicate gene promoters of the *dac* and the CG5888 genes. **(c, e)** Quantification of the *dac* PRE loop interaction scores. Hi-C interaction score in WT, ΔPRE2, or the mutant PRE lines vir, 2xPRE1, Fab-7 and en. Statistical comparisons were performed using the Kruskal–Wallis test followed by Dunn’s post-hoc pairwise test with Benjamini–Hochberg correction on all the sample considered together as in Supplementary Fig. 8c. Boxplots show median (central line), Q1=25th and Q3=75th percentiles (box limits), and Q1+1.5×IQR to Q3+1.5×IQR (whiskers), where IQR is the interquartile range.

### Loop-calling analysis

Loop-calling analysis was performed on .mcool files which were obtained and normalized via the Iterative <u>C</u>orrection and <u>E</u>igenvector decomposition algorithm (ICE) with default parameters (command “*cooler zoomify -r 400, 800, 1000, 2000, 4000, 8000, 10000, 20000, 40000 file.cool -o file.mcool --balance*”). To call loops from Micro-C maps, the following strategy as been applied. (1) Firstly, manually-curated a sets of *Gold Standard* loops per condition have been obtained from the set of loops published in Ref. (16). This step has been carried out by plotting the Micro-C maps at different resolution. (2) The *mustache* loop-caller (43) has been applied to detect loops on matrices of different resolutions with specific ranges of minimum and maximum distances between the anchors. Loop-detection has been done at resolutions 400, 800, 1000, 2000 bp between 0 and 400 kb; 800, 1000, 2000, and 4000 bp between 0 and 800 kb; 4000, 8000, 10000, and 20000 bp between 0 and 3.2 Mb, and 20000 and 40000 bp between 0 and 33 Mb. We applied *mustache* with the following *mustache* parameters *sparsityThreshold* =1.00, *iteration* = 5, and *sigmaZero* varying between 0.6 and 3.6 every 0.1. (2) Loops in the pericentromeric regions (chr2L:22113700-23513712, chr2R:1-5756000, chr3L:22933988-28110227, chr3R:1-4027467, chrX:21572709-22396687 and chrX:22929901-23542271) were filtered out because they were detected in regions with poor mappability. (3) Next, for each of the resolutions, sets of loops with False Discovery Rates (FDR) smaller than a threshold value which was set from 0.0001 to 0.1 increasing by steps of 0.0001. Loops closer than one bin were merged (command: *PyGLtools/merge.py -d resolution*). (4) To evaluate the optimal FDR per resolution, we computed how many of the Gold Standard (defined at point (1)) were detected and defined the ratio between the number of Gold Standard loops detected and the total number of loops detected (r_GS_). We finally set for each resolution the value of FDR that allows us to maximize the r_GS_. This results in a set of loops per resolution. (5) Finally, the union of all the loops that pass the FDR threshold per resolution were inspected visually. To ease this visual inspection, we plotted the entire region spanned by the loops (+/-25 bins) and marked the putative loop on the matrix using a dashed square. (6) After manual inspection, we merged the accepted loops for different resolutions where the anchors have different sizes. Each of these loops were recentered on the location within the looping area with the highest Micro-C score: the final set of loops is obtained by fixing the location of the maximum +/- 2kb. This recentering is motivated by the fact that the Shaman score remove the local genomic-distance bias and help to locate the summit of the loops. (7) In conclusion, we obtained: 358 loops from Micro-C in ED_Control, 409 loops from Micro-C in Embryo_WT, and 170 loops from Micro-C in WD_WT (**Supplementary Table 1**). Signals for H3K27me3, PH, GAF, and H3K27ac around paired loop-anchors were used to cluster loops (**Fig 1a**). Signal intensities were normalized and stratified into a defined number of clusters (k=2) using the *kmeans* function in R (version 4.4.3; www.R-project.org). Heatmaps and average signal profiles were generated using the seqplots R package (v1.23.3) (37).

### qChIP experiments

qChIP experiments were performed as described in (9) with minor modifications. Chromatin was sonicated using a Bioruptor Pico (Diagenode) for 7 min (30sec in, 30sec off). All antibodies were diluted 1/100 for the IP. After decrosslinking, DNA was purified using MicroChIP DiaPure columns from Diagenode. Enrichment of DNA fragment was analyzed by real-time PCR Light-cycler 480 (Roche). Primers used are indicated in **Supplementary Table 2**. PHO antibody is a generous gift of J. Kassis (44). Psq antibody is a generous gift from CA Berg (45). GAF, PH, E(Z) antibodies are described in (12). H3K27me3 antibody is from ActiveMotif (#39155).

## Results

### GAF binding levels correlate with chromatin looping genome-wide

To investigate the genome-wide relationship between GAF occupancy and chromatin looping, we performed CUT&RUN targeting GAF, the PRC1 subunit Polyhomeotic (PH), active and repressive histone marks (H3K27Ac and H3K27me3, respectively) alongside high-resolution Micro-C, in *Drosophila* embryos (16-20h) and in two distinct third instar larval tissues: imaginal eye and wing discs (**Supplementary Table 1**). We identified 1855, 6441 and 4272 high-confidence GAF peaks in embryos, larval imaginal eye and wing discs, respectively. Micro-C analysis revealed 409 high-confidence loops in embryos, 358 in eye imaginal discs and 170 in wing imaginal discs, with median loop sizes of 59 kb, 45 kb and 71 kb, respectively (**Supplementary Fig. 1a**). Although only a small subset of all GAF peaks engages in looping interactions (**Supplementary Fig. 1b**), GAF binding is a general hallmark of chromatin loops (**Fig. 1a**), consistent with previous reports (4,12). In embryos, 38% of loops overlap with GAF peaks, with roughly half showing GAF binding at both anchors and half at a single anchor. At the larval stage, this fraction increases to 70% in wing imaginal discs and 85% in eye imaginal discs, with most of these loops bound by GAF at both anchors (**Supplementary Fig. 1c, Supplementary** T**able 1**). These findings suggest that GAF involvement in looping becomes more prominent during development. The median sizes of GAF-overlapping loops are comparable between stages, 43 kb in embryos and 39 kb or 46 kb in eye or wing imaginal discs (**Supplementary Fig. 1a**). Importantly, chromatin loop-anchors located within repressive H3K27me3 domains exhibit a particularly strong association with GAF binding: 71%, 92% and 83% of H3K27me3-enriched loops in the embryo, eye and wing imaginal discs, respectively, overlap with GAF peaks in at least one loop anchor. These anchors are also bound by PH (**Fig. 1a**), highlighting a predominant role for GAF in mediating chromatin loops between Polycomb response elements (PREs).

Notably, GAF sites associated with loop anchors exhibited significantly higher occupancy than at non-looping sites (**Supplementary Fig. 1d**), suggesting a quantitative relationship between GAF occupancy and loop formation. To investigate this further, we ranked all GAF peaks based on their peak scores, used as a proxy for binding strength, and grouped them into quartiles where Q4 (Q1) had the highest (lowest) GAF binding strength (**Fig. 1b**). This analysis revealed a skewed distribution of GAF binding, with high-occupancy GAF peaks more likely to overlap with loop anchors (**Supplementary Fig. 1e**). Consistent with previous reports showing that GAF binds preferentially to clustered (GA)n repeats via its POZ domain (17,46), we also found that stronger GAF peaks were associated with both larger peak sizes (**Supplementary Fig. 1f**) and higher numbers of underlying GAF motifs (**Supplementary Fig. 1g,h**). To directly test whether GAF binding levels influence loop formation, we focused on pairs of GAF peaks within 500 kb of genomic separation, since the majority of GAF-associated loops fall within this range (**Supplementary Fig. 1a**). Importantly, this analysis confirmed a strong positive correlation between GAF occupancy and the likelihood of looping: the higher the GAF peak score, the greater the looping score, hence the greater the tendency for the site to engage in chromatin loops (**Fig. 1c,d**).

Together, these results demonstrate that GAF binding strength is a robust predictor of chromatin looping independently of the examined tissue and support a model in which GAF occupancy plays a critical role in establishing chromatin loops during *Drosophila* development.

Interestingly, a similar positive relationship between protein binding levels and chromatin contact frequency has previously been reported for PRC1 components, including PH (47). To explore the chromatin landscape of GAF binding sites involved in looping, we clustered CUT&RUN profiles of the H3K27ac and H3K27me3 histone marks, together with PH and chromatin accessibility (ATAC-seq) data in all GAF peaks identified in both embryos and larval imaginal eye and wing discs (**Supplementary Fig. 2**). Strikingly, the cluster with the strongest chromatin looping between GAF peaks also showed high levels of PH binding (**Supplementary Fig. 2a,b CL4**). Moreover, genomic regions that gain GAF binding at the larval stage compared to embryos display a significant increase in chromatin looping (**Supplementary Fig. 2a,b, CL3**). This increase is followed by elevated PH occupancy and enhanced chromatin accessibility, suggesting a coordinated change in chromatin state. Of note, GAF and PH co-localization and strong binding at H3K27me3 domains was associated to the strongest loops in embryos, but in larval tissues an even stronger looping was observed in a cluster of sites associated with H3K27Ac (**Supplementary Figs. 2 and 3**), suggesting that active chromatin regions might increase looping efficiency during late stages of development. In summary, the strong correlation between GAF and PH binding levels, as well as chromatin contact frequencies at all developmental stages and examined tissues, suggests that both proteins may play a functional key role in the formation of chromatin looping.

### GAF binding levels dictate PRE loop formation and PH recruitment

Since chromatin loop anchors within H3K27me3 domains show strong enrichment for both GAF and PH binding (**Fig. 1a**), and GAF sites overlapping with PH in Polycomb domains establish robust chromatin loops genome-wide (**Supplementary Fig. 3**), we next focused our analysis on dissecting the respective contributions of GAF and PH to PRE looping. Previous work demonstrated that a PRE loop at the *dac* gene locus depends on both an intact PRE sequence and GAF binding (12). Here, we found that deletion of the PRE sequence not only leads to the loss of GAF binding (12), but also to a significant reduction in PRE looping (**Supplementary Fig. 4**). To further dissect how GAF binding levels influence the formation of PRE loops, we leveraged the correlation between GAF occupancy and the number of underlying (GA)n repeat motifs (**Supplementary Fig. 1**). Therefore, we generated a series of mutant fly lines carrying targeted mutations in increasing numbers of (GA)n motifs within the *dac* PRE2 sequence (**Fig. 2a**).

qCHIP experiments confirmed that mutation of all 6 (GA)n motifs abolished both GAF and PH binding (**Fig. 2b**). Mutation of 4 (GA)n motifs resulted in a partial loss of GAF and PH binding, while mutation of only 2 (GA)n motifs led to an even lower reduction of GAF and PH levels compared to the wild type control (WT) (**Fig. 2b**). Therefore, this engineered allelic fly line series provides a quantitative gradient of GAF and PH binding levels to the PRE2 sequence. In contrast, two other PRE-bound TFs that either interact with GAF or bind to (GA)n motifs, Pleiohomeotic (PHO) and Pipsqueak (PSQ), can efficiently bind to the PRE2 sequence upon mutation of 2 or 4 (GA)n motifs, whereas their binding is only reduced when all (GA)n motifs are deleted (**Supplementary Fig. 5a**).

Next, we asked whether the strength of GAF and/or PH binding is a key determinant of PRE loop formation. To test this, we performed high resolution Micro-C experiments in the different (GA)n mutant lines and quantified the *dac* PRE loop (**Fig. 2c-d**). As previously observed, mutations of all (GA)n motifs led to a significant disruption of the PRE loop. Importantly, even mutation of just 2 (GA)n sites was sufficient to significantly reduce PRE looping, with the loss of PRE looping progressively increasing as more motifs were mutated and GAF/PH binding levels decreased (**Fig. 2c-d**). In contrast, looping frequencies between two PREs at the *NetA/B* gene locus remained unaffected by (GA)n motifs mutations at the *dac* locus, indicating that the effect is specific to the mutated PRE loop (**Fig. 2e-f**). Strikingly, the gradient of GAF and PH binding levels upon (GA)n motifs mutations closely mirrors the penetrance of a gain-of-function phenotype of the *dac* gene (**Supplementary Fig. 5b**), which has been previously shown to be induced by the specific overexpression of *dac* in the TS2 segment in pupal leg discs (13). In summary, these results indicate that high GAF binding levels are critical for robust PRE loop formation and for buffering the expression of looping-dependent mutant phenotypes at the *dac* locus, with PH binding potentially taking a part in this process.

### PH binding is dispensable for PRE looping

To directly assess whether PH is necessary for PRE loop formation, we made use of a temperature-sensitive system, previously shown to efficiently knock down PH protein levels in imaginal eye discs (38). Global loss of PH binding to chromatin also results in the strong reduction of H3K27me3 levels (38), whereas GAF binding at PREs remains unaffected (**Fig. 3a**), allowing to uncouple the respective contributions of these two chromatin factors to PRE looping. We therefore performed Micro-C experiments upon loss of PH function and quantified PRE looping at the *dac* gene locus (**Fig. 3b**). Contrary to the loss of GAF binding, PRE looping frequency between the two PREs at the *dac* gene locus was only slightly affected upon loss of PH binding (**Fig. 3c**).

To extend this analysis on a genome-wide scale, we generated aggregate plots of Micro-C signals centered on PRE loops (loops bound by PH) and compared them to loops formed in the absence of PH (non-PcG loops). Quantification of aggregated loop signals revealed a slight, though not significant, decrease in PRE looping upon loss of PH. However, a significant reduction was also observed at non-PcG loops (**Fig. 3d**), suggesting that the decrease in PRE looping is not a direct consequence of PH loss, but may instead reflect a global change in nuclear or chromosomal architecture, or a small systematic difference in the two Micro-C samples. Interestingly, although loss of PH does not specifically affect PRE looping within TADs, long range interactions between Polycomb domains are significantly reduced (**Supplementary Fig. 6**), as exemplified by the loss long range interactions between the *HOX* gene clusters (Antp-C and Bx-C, **Fig. 3e**). In contrast, long range interactions between active or null domains are not affected (**Supplementary Fig. 6**).

Altogether, these results indicate that, although PH is necessary for the compartmentalization of Polycomb domains and PH binding levels positively correlate with PRE loop strength, PH is not essential for efficient PRE looping. PH occupancy at PREs likely reflects its recruitment downstream of GAF binding and the establishment of Polycomb repression domains, rather than a direct architectural role in loop formation. In contrast, GAF, alone or in combination with other chromatin factors bound at PREs, emerges as a key driver of PRE looping.

### GAF is required for PRE looping but remains insufficient, even in conjunction with PHO binding

To investigate whether GAF binding alone is sufficient to mediate PRE looping, or whether additional PRE-binding proteins are required, we replaced the PRE2 sequence at the *dac* gene locus with a 370 bp fragment from the *Hsp26* promoter containing well-characterized (GA)n motifs (48). Notably, this sequence is known to recruit high levels of GAF without recruiting PcG proteins at its endogenous locus (**Supplementary Fig. 7a**). Importantly, we did not include the *Hsp26* TATA box to prevent artificial gene transcription (**Fig. 4a**, hsp26 mutant). qChIP analysis confirmed that GAF was efficiently recruited to the *Hsp26* sequence inserted at the *dac* TSS (**Fig. 4b**). However, neither PH nor PHO was detected at this site (**Fig. 4c,d**), indicating that GAF binding alone is not sufficient to recruit PcG proteins. The absence of PRE activity/PH recruitment of the *Hsp26* sequence was also reflected by a marked reduction in H3K27me3 levels across the *dac* domain, similar to what is observed in the PRE2 deletion line (**Supplementary Fig. 7b**). Consistent with the loss of Polycomb-mediated repression of *dac*, the Hsp26 replacement line displayed the same gain-of-function phenotype (**Supplementary Fig. 7c**) previously observed upon deletion of the PRE2 sequence (12).

To assess whether GAF binding alone, without PH, is sufficient to mediate PRE looping, we performed Hi-C experiments in third instar imaginal discs from flies carrying the Hsp26 replacement construct and compared PRE interaction frequency to the WT and the PRE2 deletion lines (**Fig. 4e**). Quantification of interactions between the PRE1 and PRE2 regions showed that the Hsp26 line exhibits significantly reduced interaction frequencies compared to WT flies, with a similar interaction frequency than the PRE2 deletion line (**Fig. 4f**). These results indicate that GAF binding alone is not sufficient to mediate PRE looping interactions within Polycomb domains, suggesting that additional PRE-associated looping factors are needed.

PHO has been proposed to be an additional looping factor, as it is preferentially enriched at loop anchors that are maintained upon loss of GAF function (15). To analyze whether the additional presence of PHO together with GAF is sufficient to mediate PRE looping, we generated a modified construct in which three PHO consensus motifs (GCCCATTT) were inserted into the *hsp26* promoter sequence (**Fig. 4a**, Hsp26+3xPHO). qChIP analysis confirmed that PHO was successfully recruited to the Hsp26+3xPHO construct, while GAF binding levels remained comparable to the unmodified Hsp26 construct (**Fig. 4d**). Importantly, despite the recruitment of both GAF and PHO, PH was still not detected at the Hsp26+3XPho construct (**Fig. 4c**), and H3K27me3 levels remained low across the *dac* domain (**Supplementary Fig. 7b**). This suggests that the simultaneous presence of GAF and PHO is not sufficient to recruit PcG proteins and establish a repressive chromatin environment at the *dac* gene locus. To evaluate the functional impact on PRE looping, we performed Hi-C experiments in the hsp26+3xPHO replacement line and quantified the interactions between PRE1 and PRE2 regions (**Fig. 4e,f**). We observed that the PRE interaction frequencies in the Hsp26+3xPHO line were significantly decreased compared to the WT, similar to the Hsp26 line (**Fig. 4f**).

These results indicate that even the combined presence of GAF and PHO is not sufficient to mediate PRE looping, arguing against the hypothesis that GAF or PHO alone are sufficient to induce looping and instead suggesting that additional factors may be required to establish PRE chromatin contacts.

### The PRE looping specificity is determined by a TF looping code

PREs act as binding platforms for a specific combination of TFs that cooperate to recruit PcG complexes and mediate PRE looping. However, only a subset of PREs form loops, and the molecular determinants of PRE-PRE looping specificity remains unclear. The specificity of PRE looping may depend on the quality and the quantity of looping factors occupying the PRE anchors, as well as the specific combination of these factors at the two looping anchors. To test whether a specific TF binding combination encoded in the PRE sequences determine PRE looping specificity, we replaced the *dac* PRE2 with different PRE sequences, namely (i) its orthologous PRE2 sequence from *Drosophila virilis* (*vir*), (ii) a homologous duplicate of the *dac* PRE1 sequence (2xPRE1), or (iii) unrelated PRE sequences from the *engrailed* (*en*) gene or the Bithorax-Complex (Fab-7) loci (**Fig. 5a**).

We first confirmed that the *vir* PRE recruits GAF and PH when inserted in the *D. melanogaster* genome (**Supplementary Fig. 8a**). Since the Fab-7 and *en* PRE sequences are present at their endogenous gene locus, we cannot directly assess the binding of these factors to the PREs when inserted at the *dac* gene locus, however both PREs are known to be bound by (high levels of) GAF and PH at their endogenous locus (9,39). Importantly, all four PRE replacements efficiently established a H3K27me3 domain and rescued the loss of H3K27me3 induced by PRE2 deletion, confirming their PRE activity (**Fig. 5a, Supplementary Fig. 8b**).

To analyze their ability to mediate PRE looping, we performed Hi-C experiments in the different PRE replacement lines (**Fig. 5b-e**). As expected, 2xPRE1 partially rescued looping caused by PRE2 deletion, although the looping frequency was slightly reduced compared to WT (**Fig. 5c**). Interestingly, the orthologous *vir* PRE also rescued looping (**Fig. 5c**). In contrast, the unrelated Fab-7 PRE, which has been shown to be able to mediate long range PRE interactions between two homologous sequences (49), showed a significantly reduced ability to loop with the *dac* PRE1 compared to the WT or the homologous and orthologous PRE sequences, with similar levels to those of the PRE2 deletion (**Fig. 5d,e**). Similarly, the *en* PRE replacement showed reduced looping frequency with the PRE1 from the *dac* locus and, although it exhibited partial recovery compared with the PRE2 deletion, this remained significantly lower than the orthologous and homologous PRE interactions (**Fig. 5d,e**). Notably, the reduction in PRE looping frequency tightly correlates with the penetrance of the previously observed *dac* gain-of-function phenotype (13), emphasizing the functional importance of this PRE loop (**Supplementary Fig. 8d**). Importantly, the *en* PRE mediates chromatin looping at its endogenous locus with the PRE from the *invected* gene, arguing against the hypothesis that PRE looping specificity is simply a matter of the quantity of looping factors bound at PREs. Instead, these results support the existence of a specific transcription factor-mediated code encoded in PRE sequences that governs the specificity of PRE-PRE looping interactions.

## Discussion

Several lines of evidence suggested a key role of GAF, either alone or in combination with PH, in mediating chromatin looping. However, due to its implication in PcG recruitment, it was so far impossible to disentangle the importance of GAF and PH for PRE looping. In this study, we investigated the direct contribution of GAF in mediating PRE loops and clarified the importance and interplay of GAF and PH for PRE looping. Furthermore, we provide a first experimental support for the existence of a TF looping code at PREs that contributes to the specificity of PRE loops.

Our main findings are summarized below: (1) A tight positive correlation throughout development and across different tissues between GAF binding levels, GAGA motif content and the ability of GAF-bound sites to mediate chromatin loops. Importantly, this correlation was demonstrated both at the genome-wide level and at a specific gene locus (*dac*). (2) *de-novo* binding of GAF and PH during development is a good predictor for the gain of chromatin looping. (3) However, PH is dispensable for PRE looping. (4) Although necessary, GAF is not sufficient to mediate chromatin loops, but requires the presence of additional looping factors, others than PH and PHO. (5) Replacement of a PRE with a homologous and/or orthologous PRE sequence can rescue PRE looping, whereas unrelated PRE sequences are significantly less effective, suggesting that PRE looping specificity is provided by two anchor sequences that recruit a specific set of compatible looping factors.

### The role of GAF in PRE looping

The implication of GAF in 3D chromatin organization has been recognized for over two decades: GAF contains a BTB/POZ oligomerization domain (17–19) that enables long-range enhancer-promoter communication and gene regulation (20,50).

More recent studies have positioned GAF as a central factor in PRE looping: (i) GAF is enriched at the loop anchors of *cis*-regulatory elements including tethering elements (TEs) (4) and PREs (12), which constitute a specific form of TE loops (13). (ii) GAF binding is necessary for both intrachromosomal PRE looping at the *dac* gene locus (13) and interchromosomal PRE interactions between homologous copies of the Fab-7 PREs (22). (iii) Mutation of GAF affects homologous PRE pairing and deletion of the POZ domain results in the loss of a subset of chromatin loops, particularly those connecting paralogous genes (15).

Here, we clarify the specific role of GAF in PRE looping. Although, *de-novo* binding of GAF during development is predictive for increased chromatin looping, our results demonstrate that GAF binding is necessary but not sufficient for loop formation. Indeed, not all loop anchors are bound by GAF, and a considerable fraction of loops involve GAF at only one anchor. This asymmetry strongly suggests that additional factors must occupy the opposite anchor to stabilize the interaction, reinforcing the idea that GAF acts in concert with other proteins rather than alone. Importantly, the loss of chromatin looping upon mutation of GAF or its binding sites likely reflects not only the loss of GAF function, but also the concomitant loss of other looping factors that either bind to the same motifs or require GAF for chromatin binding. Therefore, the previously reported essential role of GAF in chromatin looping cannot be attributed to GAF action alone, but rather reflects its functional interaction with additional looping factors that might be equally important to mediate chromatin loops.

### The role of PcG proteins and associated histone marks in PRE looping

Extensive research has linked PcG proteins, particularly PRC1 components, to PRE looping. PcG proteins are known organizers of the 3D genome architecture (26), and PRC1 is critical for both the condensation of Polycomb domains and their long-range interactions in mammals (reviewed in (6)). Moreover, PRC1 has been proposed to act as a looping factor in mammalian cells (51), potentially through oligomerization of the SAM domain of the PRC1 subunit PH. This domain is required for the condensation of Polycomb domains (23) as well as for mediating long-range interactions between them (24,25). In line with this, we observed that PH is essential for the compartmentalization of Polycomb domains during Drosophila development. Surprisingly however our results show that, although co-gain of PH and GAF binding during development is predictive of PRE looping, genome-wide loss of PH does not disrupt PRE looping. This rules out an essential role for PH in PRE loop formation and instead suggests that PH is passively recruited to PREs by GAF, without contributing directly to loop formation.

A recent study reported that changes in histone modifications, such as H3K27ac, between loop anchors/boundaries are predictive of changes in CTCF/RAD21 loops (52). Here, we found that H3K27me3, the hallmark of PcG-mediated gene silencing, does not play a major role in PRE looping, since this mark is erased upon loss of PH-binding (38), yet PRE looping is unaffected.

Together, these findings suggest that PcG proteins and their associated histone modifications are not essential for the formation of PRE loops. Instead, the roles of PRC1/PH and H3K27me3 appear to be restricted to higher-order chromatin organization, such as the interaction of Polycomb domains over long distances. In contrast, PRE loops are essentially mediated by looping factors directly present at loop anchors operating independently of PcG proteins. Therefore, these data disentangle intra-domain Polycomb-associated loop formation that do not require Polycomb proteins and associated histone marks, from long-range inter-domain Polycomb compartment interactions which strictly depend on Polycomb components.

### What might be the additional looping factors at PREs?

PREs are built up of a flexible array of DNA binding sites for a multitude of sequence-specific TFs that cooperate to recruit PcG proteins. We showed that mutating increasing numbers of (GA)n repeat motifs result in a gradual loss of GAF and PH binding levels, which negatively correlate with PRE looping efficiency. However, our subsequent finding that PH is not essential and that GAF is not sufficient for PRE looping strongly suggest that other PRE-bound proteins are involved in PRE looping, either via interaction with GAF or by directly binding to (GA)n motifs.

One candidate is PHO, which binds cooperatively with GAF at PREs (53) and is enriched at loop anchors that form independently of GAF (15). This suggests that PHO might serve as a looping factor at a subset of PREs. However, additional binding of PHO to the GAF-bound Hsp26 sequence does not restore looping, and PHO binding levels are not significantly reduced at the (GA)n_mut2 and (GA)n_mut4 PREs, although PRE looping interactions are significantly decreased. While these observations do not exclude that PHO might play a role in PRE looping, they suggest that other factors must be important.

Pipsqueak (PSQ) is another candidate, as it binds to (GA)n motifs at PRE sequences, interacts with GAF (54) and possesses a BTB-POZ domain that is able to oligomerize. In addition, a specific isoform of PSQ has been implicated in chromatin looping (55). However, like PHO, PSQ binding to the (GA)n_mut2 and (GA)n_mut4 PREs is not reduced, again indicating that loss of looping is not due to reduced PSQ binding, and suggesting instead that additional TFs/looping factors are required.

Other proteins that may contribute include CLAMP, a (GA)n-binding protein implicated in long-range chromatin interactions (56), as well as other BTB/POZ domain-containing proteins that interact with GAF, such as LOLAL (57) or BAB1/2 (58), or the recently characterized Vostok protein, which functionally cooperates with GAF in forming a subset of loops in the *Drosophila* brain (59).

In summary, our findings argue that, similar to PcG recruitment to PREs, which involves the combination of many TFs, PRE looping is not governed by a single chromatin factor, but rather depends on the specific combinatorial and quantitative interplay of multiple looping factors at the loop anchors.

### Specificity of PRE loops

The observation that only a subset of PREs form loops raises a fundamental question: what determines the specificity to these chromatin interactions? Importantly, the transcriptional state of the underlying genes does not appear to be a defining factor, as PRE loops occur independently of whether the associated locus is active or repressed (13,14). Conversely, several lines of evidence suggest that the sequence content of PREs plays a major role. For instance, sequence homology between PREs has been shown to be important for long-range PRE interactions (49), and many genes associated with PRE loop anchors are paralogous-derived from gene duplications-or are functionally related (15,16).

However, not all looping interactions can be explained by sequence homology alone. The PRE loop at the *dac* locus, for example, occurs between two non-homologous sequences associated with unrelated genes (*dac* and *CG5888*). Moreover, the *D. virilis* ortholog of PRE2 can robustly rescue PRE2 function and restore looping with PRE1 at the *dac* locus, although no significant sequence similarity is found when blasting the two orthologous sequences. Finally, prior work has shown that while Polycomb domains, PRC1 binding and TF binding are a well conserved feature of the genome across *Drosophila* species, the underlying PRE DNA sequences evolve rapidly (39). These findings suggest that it is not primary sequence similarity *per se*, but rather a conserved combinatorial logic of TF binding sites that ensures PRE compatibility.

Thus, the specificity of PRE loops can be explained by two mutually non-exclusive hypotheses: first, looping may depend on reaching a threshold number of looping factors bound at both PRE anchors, such that PREs capable of looping have accumulated sufficient total binding. Supporting this idea, we observed a positive correlation between GAF binding strength and PRE looping efficiency. Additionally, PREs might contain different combinations of binding sites for TFs, in particular those containing POZ domains (60), and those with the closest match interact with one another to form a specific regulatory loop. Our PRE replacement results are in favor of this hypothesis, as replacing a PRE with a homologous sequence is significantly more efficient in mediating looping interactions compared to unrelated PRE sequences, even when they all recruit high levels of looping factors. This supports the existence of a “TF looping code” that guides PRE-PRE recognition and chromatin looping.

In conclusion, our study clarifies the importance of GAF in PRE looping and reveals that, contrary to prior assumptions, PH is dispensable for local PRE loops. We provide the first experimental evidence for a TF looping code underlying PRE loop specificity. Given the fact that replacing the *dac* PRE2 with heterologous PREs generally does not fully rescue the *dac* phenotype observed upon deletion of *dac* PRE2, we hypothesize that PRE specificity does play an important regulatory role in the genome, which might in part depend on their ability to set up higher-order 3D chromatin interactions. Future work will be required to decode this combinatorial molecular logic governing PRE looping specificity.

## Data Availability

All sequencing data are available at GEO under the series: GSE247377 (CUT&RUN); GSE310299 (Micro-C); GSE310296 (Hi-C). All original codes were deposited on GitHub (https://github.com/cavallifly/Sabaris_et_al_2025) and are publicly available as of the date of publication. Any additional information required to reanalyze the data reported in this paper is available from the corresponding author upon request. Fly lines are available on request.

## Acknowledgments

We are grateful to J. Kassis, for generous gift of PHO and antibodies, and to CA Berg for PSQ antibodies. We would like to thank MRI and Drosophila facilities (BioCampus Montpellier), CNRS, INSERM, and University de Montpellier.

## Author Contributions

G.C., G.S. and B.S. conceived this study. B.S. generated mutant fly lines. G.S., S.D., A.M.P. and B.S. performed experiments. G.S., L.F. and B.S. performed CUT&RUN and Micro-C experiments. G.S. performed computational analysis of CUT&RUN experiments. M.D.S., G.S. and G.P. performed bioinformatic analysis of Micro-C and Hi-C experiments. B.S., G.S. and G.C. wrote the manuscript.

## References

1. Nora, E.P., Lajoie, B.R., Schulz, E.G., Giorgetti, L., Okamoto, I., Servant, N., Piolot, T., van Berkum, N.L., Meisig, J., Sedat, J., et al. (2012) Spatial partitioning of the regulatory landscape of the X-inactivation centre. Nature, 485, 381–385.

2. Sexton, T., Yaffe, E., Kenigsberg, E., Bantignies, F., Leblanc, B., Hoichman, M., Parrinello, H., Tanay, A. and Cavalli, G. (2012) Three-dimensional folding and functional organization principles of the Drosophila genome. Cell, 148, 458–472.

3. Dixon, J.R., Selvaraj, S., Yue, F., Kim, A., Li, Y., Shen, Y., Hu, M., Liu, J.S. and Ren, B. (2012) Topological domains in mammalian genomes identified by analysis of chromatin interactions. Nature, 485, 376–380.

4. Batut, P.J., Bing, X.Y., Sisco, Z., Raimundo, J., Levo, M. and Levine, M.S. (2022) Genome organization controls transcriptional dynamics during development. Science, 375, 566–570.

5. Schuettengruber, B. and Cavalli, G. (2009) Recruitment of polycomb group complexes and their role in the dynamic regulation of cell fate choice. Development, 136, 3531–3542.

6. Schuettengruber, B., Bourbon, H.M., Di Croce, L. and Cavalli, G. (2017) Genome Regulation by Polycomb and Trithorax: 70 Years and Counting. Cell, 171, 34–57.

7. Denaud, S., Sabaris, G., Di Stefano, M., Papadopoulos, G.L., Schuettengruber, B. and Cavalli, G. (2025) Determining the functional relationship between epigenetic and physical chromatin domains in Drosophila. Genome Biol, 26, 116.

8. Schwartz, Y.B., Kahn, T.G., Nix, D.A., Li, X.Y., Bourgon, R., Biggin, M. and Pirrotta, V. (2006) Genome-wide analysis of Polycomb targets in Drosophila melanogaster. Nat Genet, 38, 700–705.

9. Schuettengruber, B., Ganapathi, M., Leblanc, B., Portoso, M., Jaschek, R., Tolhuis, B., van Lohuizen, M., Tanay, A. and Cavalli, G. (2009) Functional anatomy of polycomb and trithorax chromatin landscapes in Drosophila embryos. PLoS Biol, 7, e13.

10. Eagen, K.P., Aiden, E.L. and Kornberg, R.D. (2017) Polycomb-mediated chromatin loops revealed by a subkilobase-resolution chromatin interaction map. Proc Natl Acad Sci U S A, 114, 8764–8769.

11. Hug, C.B., Grimaldi, A.G., Kruse, K. and Vaquerizas, J.M. (2017) Chromatin Architecture Emerges during Zygotic Genome Activation Independent of Transcription. Cell, 169, 216–228 e219.

12. Ogiyama, Y., Schuettengruber, B., Papadopoulos, G.L., Chang, J.M. and Cavalli, G. (2018) Polycomb-Dependent Chromatin Looping Contributes to Gene Silencing during Drosophila Development. Mol Cell, 71, 73–88 e75.

13. Denaud, S., Bardou, M., Papadopoulos, G.L., Grob, S., Di Stefano, M., Sabaris, G., Nollmann, M., Schuettengruber, B. and Cavalli, G. (2024) A PRE loop at the dac locus acts as a topological chromatin structure that restricts and specifies enhancer-promoter communication. Nat Struct Mol Biol.

14. Brown, J.L., Zhang, L., Rocha, P.P., Kassis, J.A. and Sun, M.A. (2024) Polycomb protein binding and looping in the ON transcriptional state. Sci Adv, 10, eadn1837.

15. Li, X., Tang, X., Bing, X., Catalano, C., Li, T., Dolsten, G., Wu, C. and Levine, M. (2023) GAGA-associated factor fosters loop formation in the Drosophila genome. Mol Cell.

16. Pollex, T., Marco-Ferreres, R., Ciglar, L., Ghavi-Helm, Y., Rabinowitz, A., Viales, R.R., Schaub, C., Jankowski, A., Girardot, C. and Furlong, E.E.M. (2024) Chromatin gene-gene loops support the cross-regulation of genes with related function. Mol Cell, 84, 822–838 e828.

17. Katsani, K.R., Hajibagheri, M.A. and Verrijzer, C.P. (1999) Co-operative DNA binding by GAGA transcription factor requires the conserved BTB/POZ domain and reorganizes promoter topology. EMBO J, 18, 698–708.

18. Bardwell, V.J. and Treisman, R. (1994) The POZ domain: a conserved protein-protein interaction motif. Genes Dev, 8, 1664–1677.

19. Espinas, M.L., Jimenez-Garcia, E., Vaquero, A., Canudas, S., Bernues, J. and Azorin, F. (1999) The N-terminal POZ domain of GAGA mediates the formation of oligomers that bind DNA with high affinity and specificity. J Biol Chem, 274, 16461–16469.

20. Mahmoudi, T., Katsani, K.R. and Verrijzer, C.P. (2002) GAGA can mediate enhancer function in trans by linking two separate DNA molecules. EMBO J, 21, 1775–1781.

21. Mohana, G., Dorier, J., Li, X., Mouginot, M., Smith, R.C., Malek, H., Leleu, M., Rodriguez, D., Khadka, J., Rosa, P. et al. (2023) Chromosome-level organization of the regulatory genome in the Drosophila nervous system. Cell, 186, 3826–3844 e3826.

22. Fitz-James, M.H., Sabaris, G., Sarkies, P., Bantignies, F. and Cavalli, G. (2025) Interchromosomal contacts between regulatory regions trigger stable transgenerational epigenetic inheritance in Drosophila. Mol Cell, 85, 677–691 e676.

23. Kundu, S., Ji, F., Sunwoo, H., Jain, G., Lee, J.T., Sadreyev, R.I., Dekker, J. and Kingston, R.E. (2017) Polycomb Repressive Complex 1 Generates Discrete Compacted Domains that Change during Differentiation. Mol Cell, 65, 432–446 e435.

24. Isono, K., Endo, T.A., Ku, M., Yamada, D., Suzuki, R., Sharif, J., Ishikura, T., Toyoda, T., Bernstein, B.E. and Koseki, H. (2013) SAM domain polymerization links subnuclear clustering of PRC1 to gene silencing. Dev Cell, 26, 565–577.

25. Wani, A.H., Boettiger, A.N., Schorderet, P., Ergun, A., Munger, C., Sadreyev, R.I., Zhuang, X., Kingston, R.E. and Francis, N.J. (2016) Chromatin topology is coupled to Polycomb group protein subnuclear organization. Nat Commun, 7, 10291.

26. Boyle, S., Flyamer, I.M., Williamson, I., Sengupta, D., Bickmore, W.A. and Illingworth, R.S. (2020) A central role for canonical PRC1 in shaping the 3D nuclear landscape. Genes Dev, 34, 931–949.

27. Chetverina, D., Erokhin, M. and Schedl, P. (2021) GAGA factor: a multifunctional pioneering chromatin protein. Cell Mol Life Sci, 78, 4125–4141.

28. Sabaris, G., Schuettengruber, B., Papadopoulos, G.L., Coronado-Zamora, M., Fitz-James, M.H., Gonzalez, J. and Cavalli, G. (2025) A mechanistic basis for genetic assimilation in natural fly populations. Proc Natl Acad Sci U S A, 122, e2415982122.

29. Langmead, B., Trapnell, C., Pop, M. and Salzberg, S.L. (2009) Ultrafast and memory-efficient alignment of short DNA sequences to the human genome. Genome Biol, 10, R25.

30. Li, H., Handsaker, B., Wysoker, A., Fennell, T., Ruan, J., Homer, N., Marth, G., Abecasis, G., Durbin, R. and Genome Project Data Processing, S. (2009) The Sequence Alignment/Map format and SAMtools. Bioinformatics, 25, 2078-2079.

31. Tarasov, A., Vilella, A.J., Cuppen, E., Nijman, I.J. and Prins, P. (2015) Sambamba: fast processing of NGS alignment formats. Bioinformatics, 31, 2032–2034.

32. Zhang, Y., Liu, T., Meyer, C.A., Eeckhoute, J., Johnson, D.S., Bernstein, B.E., Nusbaum, C., Myers, R.M., Brown, M., Li, W. et al. (2008) Model-based analysis of ChIP-Seq (MACS). Genome Biol, 9, R137.

33. Lawrence, M., Huber, W., Pages, H., Aboyoun, P., Carlson, M., Gentleman, R., Morgan, M.T. and Carey, V.J. (2013) Software for computing and annotating genomic ranges. PLoS Comput Biol, 9, e1003118.

34. Ramirez, F., Ryan, D.P., Gruning, B., Bhardwaj, V., Kilpert, F., Richter, A.S., Heyne, S., Dundar, F. and Manke, T. (2016) deepTools2: a next generation web server for deep-sequencing data analysis. Nucleic Acids Res, 44, W160–165.

35. Lopez-Delisle, L., Rabbani, L., Wolff, J., Bhardwaj, V., Backofen, R., Gruning, B., Ramirez, F. and Manke, T. (2021) pyGenomeTracks: reproducible plots for multivariate genomic datasets. Bioinformatics, 37, 422–423.

36. Grant, C.E., Bailey, T.L. and Noble, W.S. (2011) FIMO: scanning for occurrences of a given motif. Bioinformatics, 27, 1017–1018.

37. Stempor, P. and Ahringer, J. (2016) SeqPlots - Interactive software for exploratory data analyses, pattern discovery and visualization in genomics. Wellcome Open Res, 1, 14.

38. Parreno, V., Loubiere, V., Schuettengruber, B., Fritsch, L., Rawal, C.C., Erokhin, M., Gyorffy, B., Normanno, D., Di Stefano, M., Moreaux, J., et al. (2024) Transient loss of Polycomb components induces an epigenetic cancer fate. Nature, 629, 688–696.

39. Schuettengruber, B., Oded Elkayam, N., Sexton, T., Entrevan, M., Stern, S., Thomas, A., Yaffe, E., Parrinello, H., Tanay, A. and Cavalli, G. (2014) Cooperativity, specificity, and evolutionary stability of Polycomb targeting in Drosophila. Cell Rep, 9, 219–233.

40. Ramalingam, V., Natarajan, M., Johnston, J. and Zeitlinger, J. (2021) TATA and paused promoters active in differentiated tissues have distinct expression characteristics. Mol Syst Biol, 17, e9866.

41. Vizcaya-Molina, E., Klein, C.C., Serras, F., Mishra, R.K., Guigo, R. and Corominas, M. (2018) Damage-responsive elements in Drosophila regeneration. Genome Res, 28, 1852–1866.

42. Olivares-Chauvet, P., Mukamel, Z., Lifshitz, A., Schwartzman, O., Elkayam, N.O., Lubling, Y., Deikus, G., Sebra, R.P. and Tanay, A. (2016) Capturing pairwise and multi-way chromosomal conformations using chromosomal walks. Nature, 540, 296–300.

43. Roayaei Ardakany, A., Gezer, H.T., Lonardi, S. and Ay, F. (2020) Mustache: multi-scale detection of chromatin loops from Hi-C and Micro-C maps using scale-space representation. Genome Biol, 21, 256.

44. Brown, J.L., Fritsch, C., Mueller, J. and Kassis, J.A. (2003) The Drosophila pho-like gene encodes a YY1-related DNA binding protein that is redundant with pleiohomeotic in homeotic gene silencing. Development, 130, 285–294.

45. Horowitz, H. and Berg, C.A. (1996) The Drosophila pipsqueak gene encodes a nuclear BTB-domain-containing protein required early in oogenesis. Development, 122, 1859–1871.

46. van Steensel, B., Delrow, J. and Bussemaker, H.J. (2003) Genomewide analysis of Drosophila GAGA factor target genes reveals context-dependent DNA binding. Proc Natl Acad Sci U S A, 100, 2580–2585.

47. Loubiere, V., Papadopoulos, G.L., Szabo, Q., Martinez, A.M. and Cavalli, G. (2020) Widespread activation of developmental gene expression characterized by PRC1-dependent chromatin looping. Sci Adv, 6, eaax4001.

48. Sandaltzopoulos, R., Mitchelmore, C., Bonte, E., Wall, G. and Becker, P.B. (1995) Dual regulation of the Drosophila hsp26 promoter in vitro. Nucleic Acids Res, 23, 2479–2487.

49. Bantignies, F., Grimaud, C., Lavrov, S., Gabut, M. and Cavalli, G. (2003) Inheritance of Polycomb-dependent chromosomal interactions in Drosophila. Genes Dev, 17, 2406–2420.

50. Petrascheck, M., Escher, D., Mahmoudi, T., Verrijzer, C.P., Schaffner, W. and Barberis, A. (2005) DNA looping induced by a transcriptional enhancer in vivo. Nucleic Acids Res, 33, 3743–3750.

51. Crispatzu, G., Rehimi, R., Pachano, T., Bleckwehl, T., Cruz-Molina, S., Xiao, C., Mahabir, E., Bazzi, H. and Rada-Iglesias, A. (2021) The chromatin, topological and regulatory properties of pluripotency-associated poised enhancers are conserved in vivo. Nat Commun, 12, 4344.

52. Bond, M.L., Davis, E.S., Quiroga, I.Y., Dey, A., Kiran, M., Love, M.I., Won, H. and Phanstiel, D.H. (2023) Chromatin loop dynamics during cellular differentiation are associated with changes to both anchor and internal regulatory features. Genome Res, 33, 1258–1268.

53. Mahmoudi, T., Zuijderduijn, L.M., Mohd-Sarip, A. and Verrijzer, C.P. (2003) GAGA facilitates binding of Pleiohomeotic to a chromatinized Polycomb response element. Nucleic Acids Res, 31, 4147–4156.

54. Schwendemann, A. and Lehmann, M. (2002) Pipsqueak and GAGA factor act in concert as partners at homeotic and many other loci. Proc Natl Acad Sci U S A, 99, 12883–12888.

55. Gutierrez-Perez, I., Rowley, M.J., Lyu, X., Valadez-Graham, V., Vallejo, D.M., Ballesta-Illan, E., Lopez-Atalaya, J.P., Kremsky, I., Caparros, E., Corces, V.G. et al. (2019) Ecdysone-Induced 3D Chromatin Reorganization Involves Active Enhancers Bound by Pipsqueak and Polycomb. Cell Rep, 28, 2715–2727 e2715.

56. Jordan, W., 3rd and Larschan, E. (2021) The zinc finger protein CLAMP promotes long-range chromatin interactions that mediate dosage compensation of the Drosophila male X-chromosome. Epigenetics Chromatin, 14, 29.

57. Faucheux, M., Roignant, J.Y., Netter, S., Charollais, J., Antoniewski, C. and Theodore, L. (2003) batman Interacts with polycomb and trithorax group genes and encodes a BTB/POZ protein that is included in a complex containing GAGA factor. Mol Cell Biol, 23, 1181–1195.

58. Bonchuk, A., Denisov, S., Georgiev, P. and Maksimenko, O. (2011) Drosophila BTB/POZ domains of “ttk group” can form multimers and selectively interact with each other. J Mol Biol, 412, 423–436.

59. Hu, J., Li, X., Lomaev, D., Vorobyeva, N.E., Levine, M., Erokhin, M. and Chetverina, D. (2025) Vostok: A looping factor for the organization of the regulatory genome in the Drosophila brain. Mol Cell, 85, 2442-2451 e2445.

60. Li, X. and Levine, M. (2024) What are tethering elements? Curr Opin Genet Dev, 84, 102151.

