## Supplementary Figures for "Looping specificity of Polycomb response elements requires GAF and a combinatorial code of looping factors"

### Figure S1

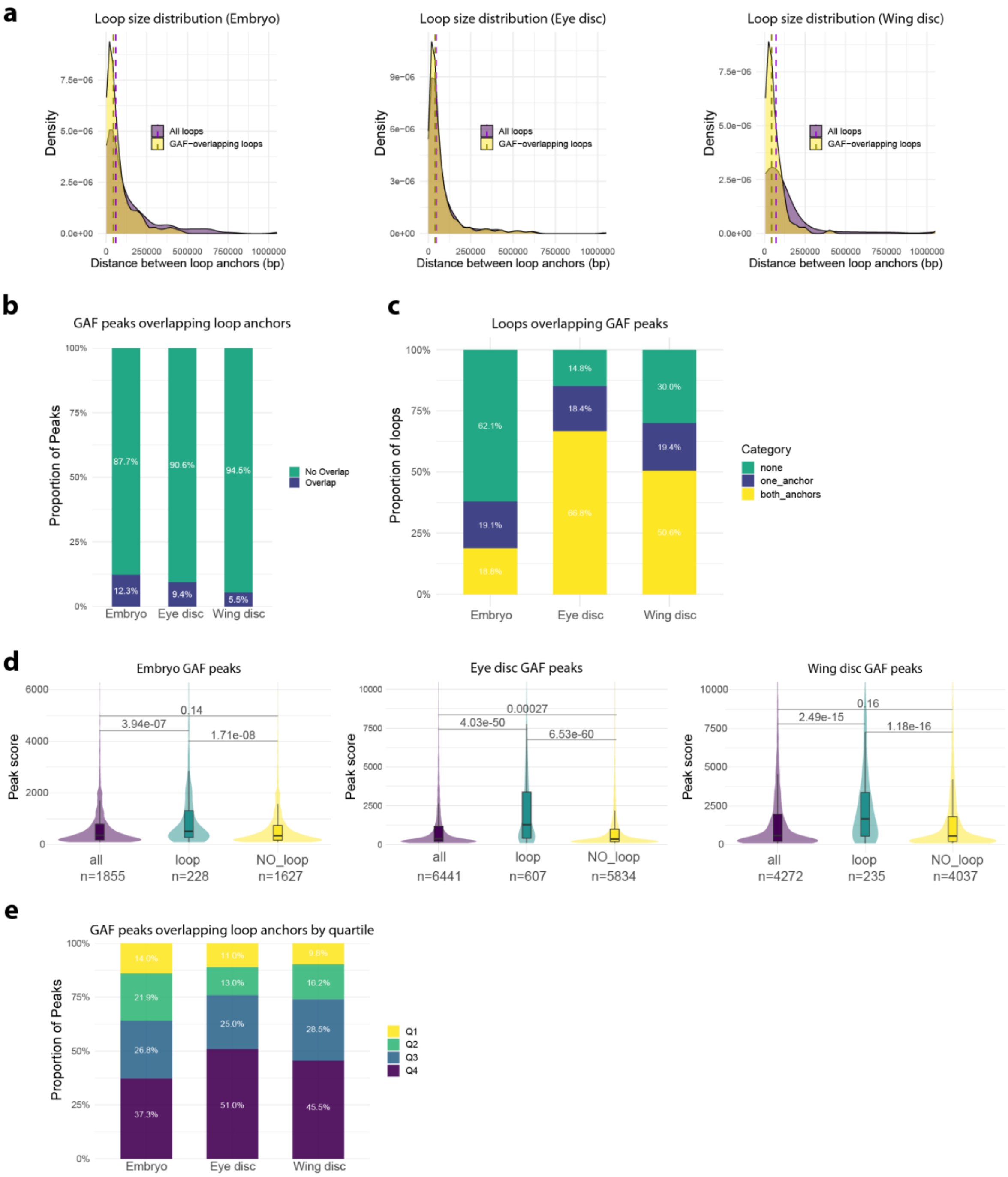
a
Loop size distribution (Embryo)
Loop size distribution (Eye disc)
Loop size distribution (Wing disc)
Density
All loops
GAF−overlapping loops
Distance between loop anchors (bp)
b
GAF peaks overlapping loop anchors
Proportion of Peaks
No Overlap
Overlap
87.7%
90.6%
94.5%
12.3%
9.4%
5.5%
Embryo
Eye disc
Wing disc
c
Loops overlapping GAF peaks
Proportion of loops
Category
none
one_anchor
both_anchors
62.1%
19.1%
18.8%
14.8%
18.4%
66.8%
30.0%
19.4%
50.6%
d
Embryo GAF peaks
Eye disc GAF peaks
Wing disc GAF peaks
Peak score
0.14
3.94e-07
1.71e-08
0.00027
4.03e-50
6.53e-60
0.16
2.49e-15
1.18e-16
all
loop
NO_loop
n=1855
n=228
n=1627
n=6441
n=607
n=5834
n=4272
n=235
n=4037
e
GAF peaks overlapping loop anchors by quartile
Proportion of Peaks
Q1
Q2
Q3
Q4
14.0%
21.9%
26.8%
37.3%
11.0%
13.0%
25.0%
51.0%
9.8%
16.2%
28.5%
45.5%

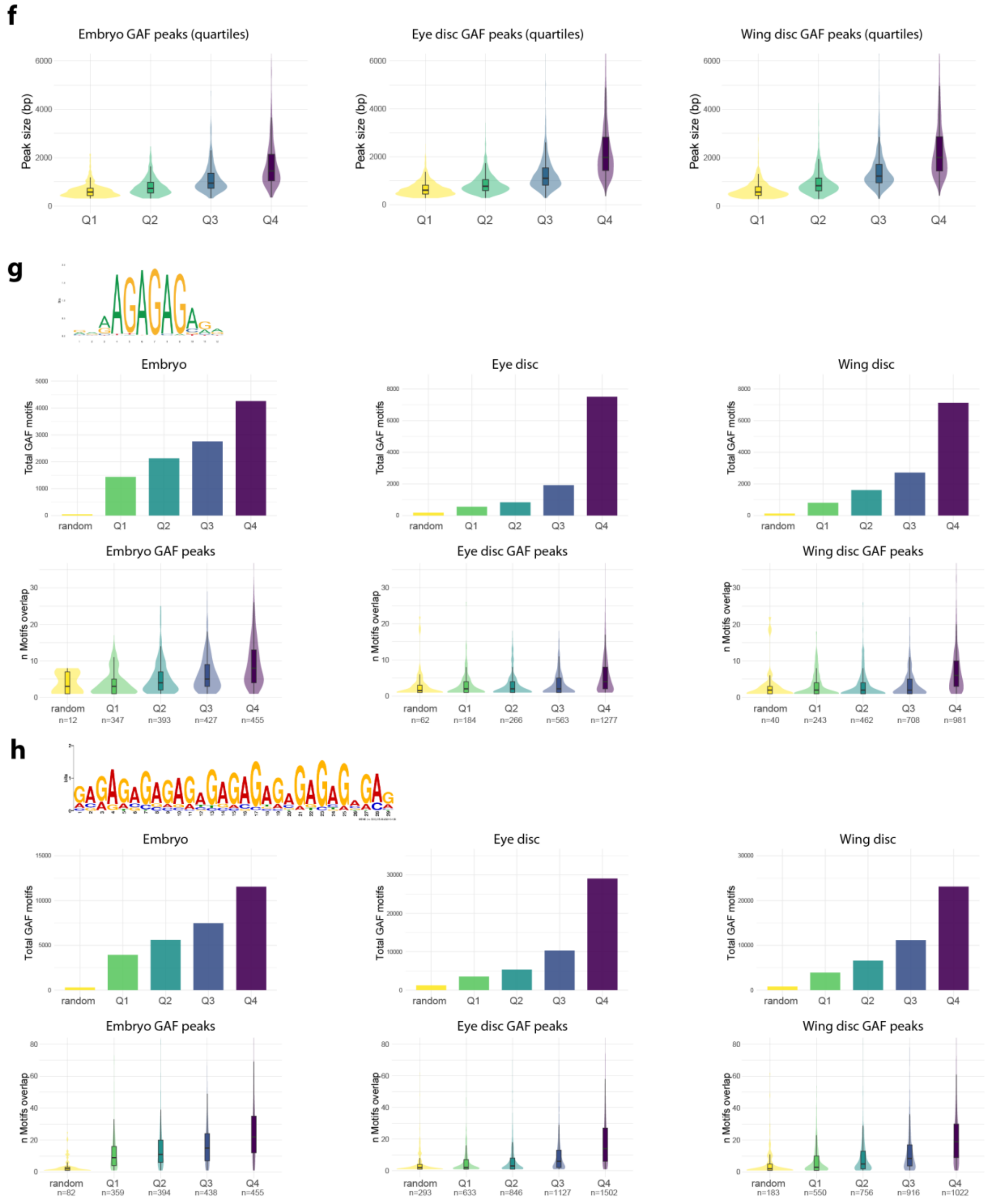


**Supplementary Figure 1: Characterization of GAF peaks that form chromatin looping**
**(a)** Histograms showing the distribution of distances between loop anchors for all loops and for GAF-overlapping loops (at least one anchor bound by GAF) in embryos and larval eye or wing imaginal discs. **(b)** Percentages of GAF peaks that overlap with loop anchors in embryos and larval eye or wing imaginal discs. **(c)** Percentages of loops that either overlap, in at least one anchor or both anchors, or do not overlap with GAF peaks in embryos and larval eye or wing imaginal discs. **(d)** Violin plots showing MACS3 peak score distributions for all GAF peaks (all), in peaks that overlap with loop anchors (loop) and in peaks that do not loop (NO_loop), in embryos and larval eye or wing imaginal discs. Statistical significance of median differences was assessed using Dunn's test with Benjamini-Hochberg correction. **(e)** Proportion of GAF peaks overlapping with loop anchors, ranked by MACS3 peak score quartiles in embryos and larval eye or wing imaginal discs. **(f)** Violin plots showing the distribution of peak sizes across GAF peak quartiles in embryos and larval eye or wing imaginal discs**. (g-h)** Total number (top) and number of motif-matches per peak (bottom) using either the canonical GAF motif (MA0205.2) **(g)** or the GA29 MEME-1 motif **(h)** in random peaks and GAF peak quartiles from embryos and larval eye or wing imaginal discs.

### Figure S2

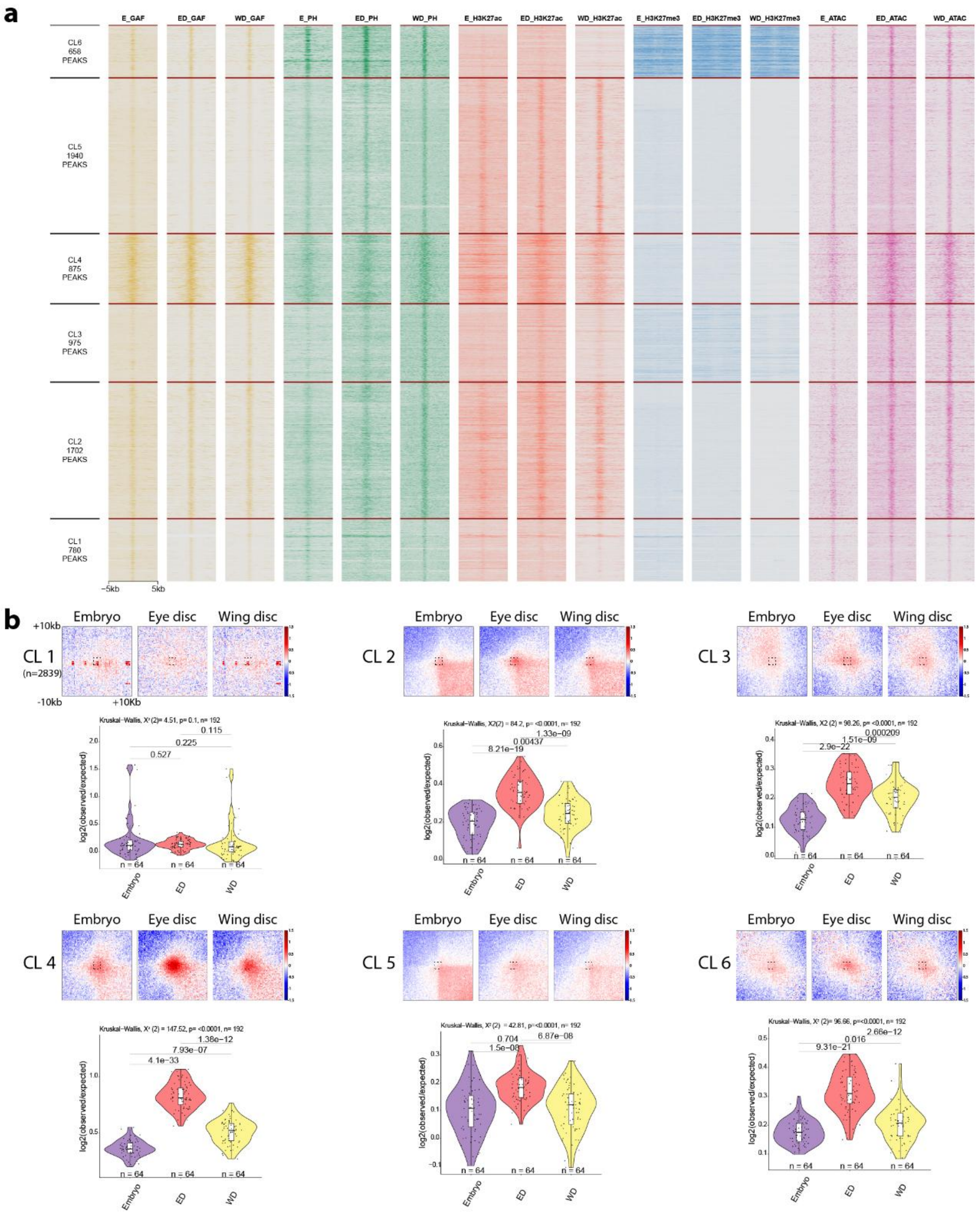


**Supplementary Figure 2: GAF binding dynamics correlate with chromatin looping**

**(a)** Heatmap showing the *k*-means clustering of GAF, PH, H3K27ac, H3K27me3, and ATAC-seq coverage around all merged GAF peaks from embryos and larval eye and wing imaginal discs (E: embryo; ED: eye disc; WD: wing disc). **(b)** Aggregate Micro-C map in embryos and larval eye and wing imaginal discs at 250 bp resolution centered at GAF peaks clustered as in panel **(a)** at distance separation between 10 and 500 kb and quantification of the signal in the central 8x8 square. Statistical comparisons were performed using the Kruskal–Wallis test followed by Dunn's post-hoc pairwise test with Benjamini–Hochberg correction. Boxplots show median (central line), Q1=25th and Q3=75th percentiles (box limits), and Q1+1.5×IQR to Q3+1.5×IQR (whiskers), where IQR is the interquartile range.

### Figure S3

**a**

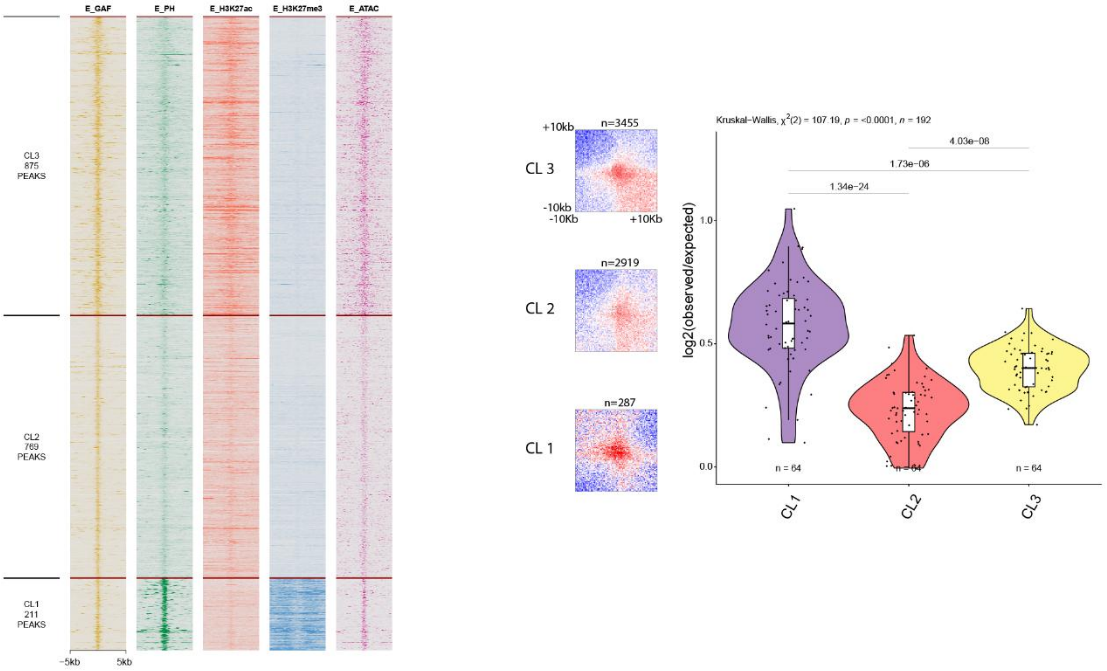
E_GAF
E_PH
E_H3K27ac
E_H3K27me3
E_ATAC
CL3
875
PEAKS
CL2
769
PEAKS
CL1
211
PEAKS
−5kb
5kb
+10kb
n=3455
CL 3
-10kb
-10Kb
+10Kb
n=2919
CL 2
n=287
CL 1
Kruskal−Wallis, $\chi^2(2)$ = 107.19, $p$ = <0.0001, $n$ = 192
4.03e−08
1.73e−06
1.34e−24
log2(observed/expected)
n = 64
n = 64
n = 64
CL1
CL2
CL3

**b**

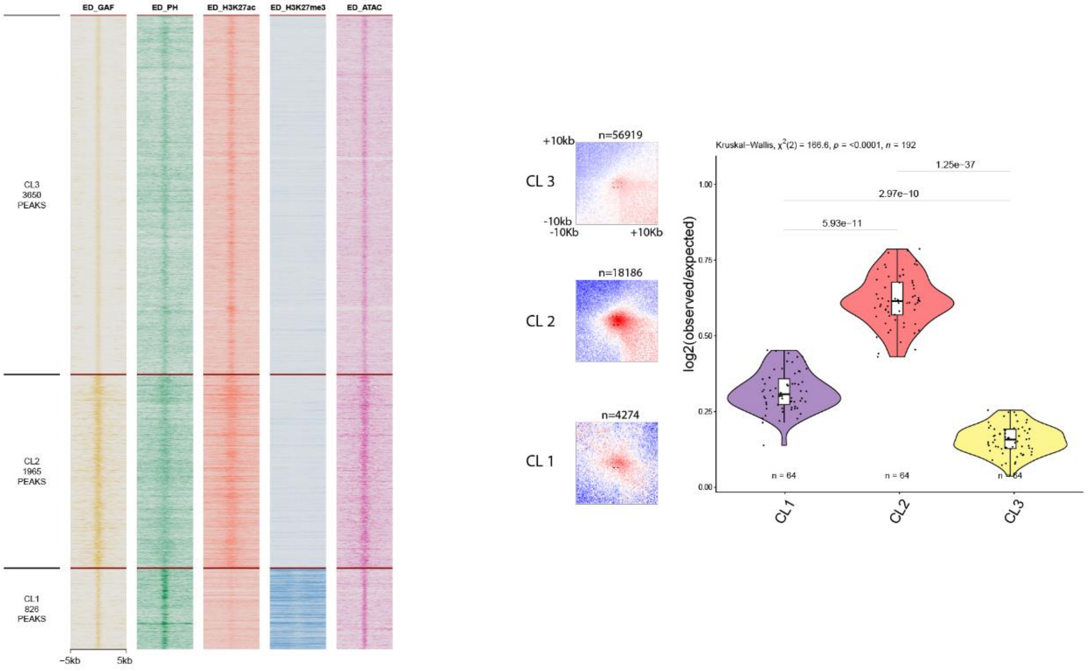
ED_GAF
ED_PH
ED_H3K27ac
ED_H3K27me3
ED_ATAC
CL3
3650
PEAKS
CL2
1965
PEAKS
CL1
826
PEAKS
−5kb
5kb
+10kb
n=56919
CL 3
-10kb
-10Kb
+10Kb
n=18186
CL 2
n=4274
CL 1
Kruskal−Wallis, $\chi^2(2)$ = 166.6, $p$ = <0.0001, $n$ = 192
1.25e−37
2.97e−10
5.93e−11
log2(observed/expected)
n = 64
n = 64
n = 64
CL1
CL2
CL3

c

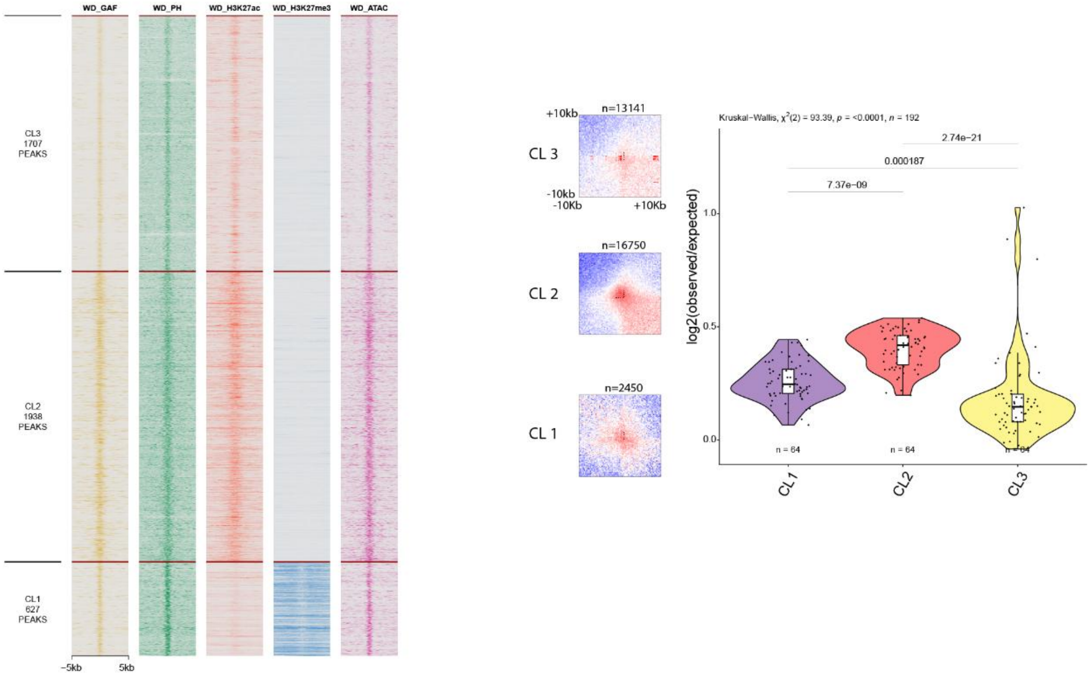


**Supplementary Figure 3: PREs bound by GAF form strong loop interactions**

**(a-c)** Heatmaps showing *k*-means clustering of GAF, PH, H3K27ac, H3K27me3, and ATAC-seq coverage around GAF peaks in embryos **(a)** and larval eye **(b)** or wing imaginal discs **(c)**. Aggregate Micro-C map at 250 bp resolution centered GAF peaks, which are clustered as in the corresponding left panels at distance separation between 10 and 500 kb and quantification of the signal in the central 8x8 square. Statistical comparisons were performed using the Kruskal–Wallis test followed by Dunn's post-hoc pairwise test with Benjamini–Hochberg correction. Boxplots show median (central line), Q1=25th and Q3=75th percentiles (box limits), and Q1+1.5×IQR to Q3+1.5×IQR (whiskers), where IQR is the interquartile range.

#### Figure S4

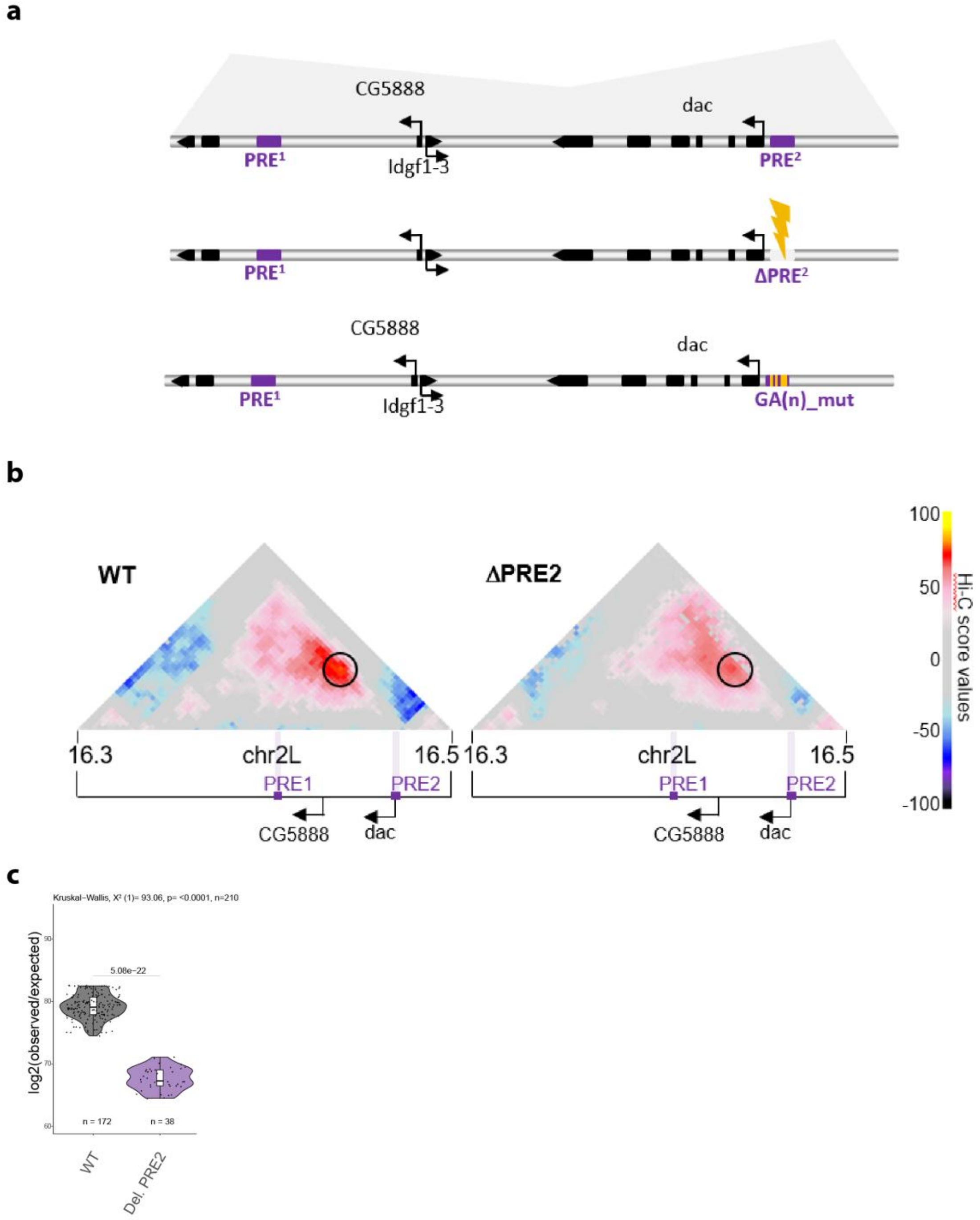


**Supplementary Figure 4: Mutation of PRE2 sequence results in loss of PRE looping**
**(a)** Schematic representation of the *dac* gene locus and mutated PRE2 lines. **(b)** Hi-C score (**Material and Methods**) maps of a 200 kb region at 3kb resolution on chromosome 2L at *dac* gene locus in imaginal discs in WT or ΔPRE2 flies. Black circle indicates the position of the *dac* PRE loop. Violet bars indicate position of PREs. Black arrows indicate promoters of the *dac* and the *CG5888* genes. **(c)** Quantification of the *dac* PRE loop interaction scores. Hi-C interaction score in WT and ΔPRE2 flies. The number of points per distribution is reported (**Material and Methods**). Boxplots show median (central line), Q1=25th and Q3=75th percentiles (box limits), and Q1+1.5×IQR to Q3+1.5×IQR (whiskers), where IQR is the interquartile range. Statistical comparisons were performed using the Kruskal–Wallis test followed by Dunn's post-hoc pairwise test with Benjamini–Hochberg correction on all the sample considered together as in **Supplementary Fig. 8c**.

**Figure S5**

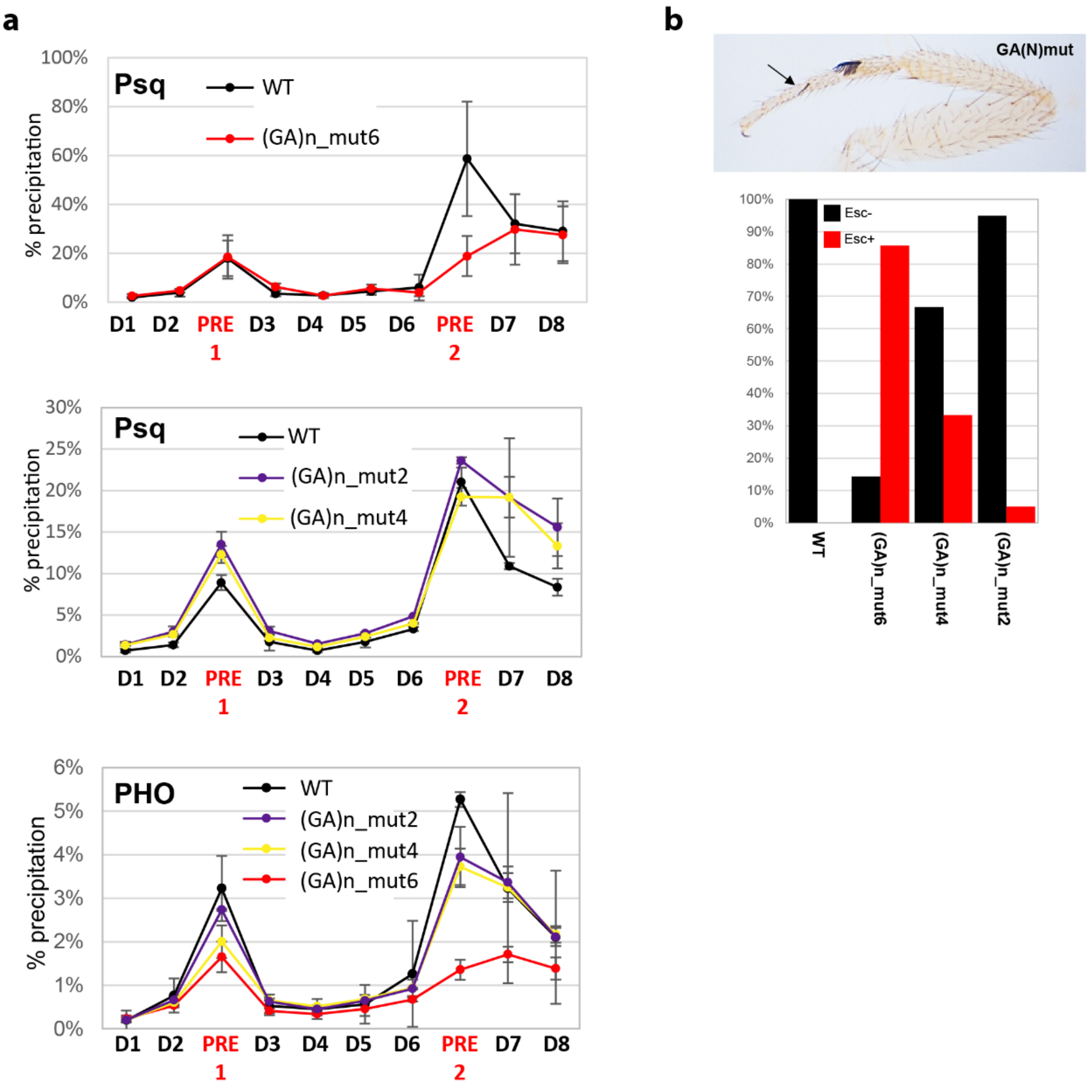


**Supplementary Figure 5: Consequences of mutating different numbers of GAGA motifs on TF binding**
**(a)** qChIP experiments using PSQ and PHO antibodies in WT, (GA)n_mut2, (GA)n_mut4 and (GA)n_mut6 fly lines. D1-D8, and PRE1, 2 indicate PCR amplicons used for qChIP experiments along the *dac* gene locus (**Supplementary Table 2**). The housekeeping gene *Rp49* was used as negative control, the *engrailed* PRE as a positive control. Data are presented as the mean values ± s.d (error bars) of two independent replicates. **(b)** Quantification of the extra sex comb (ESC) phenotype in the indicated fly lines grown at 28°C. A minimum of 50 male flies was scored. Above a representative example of a mutant phenotype leg is shown.

#### Figure S6

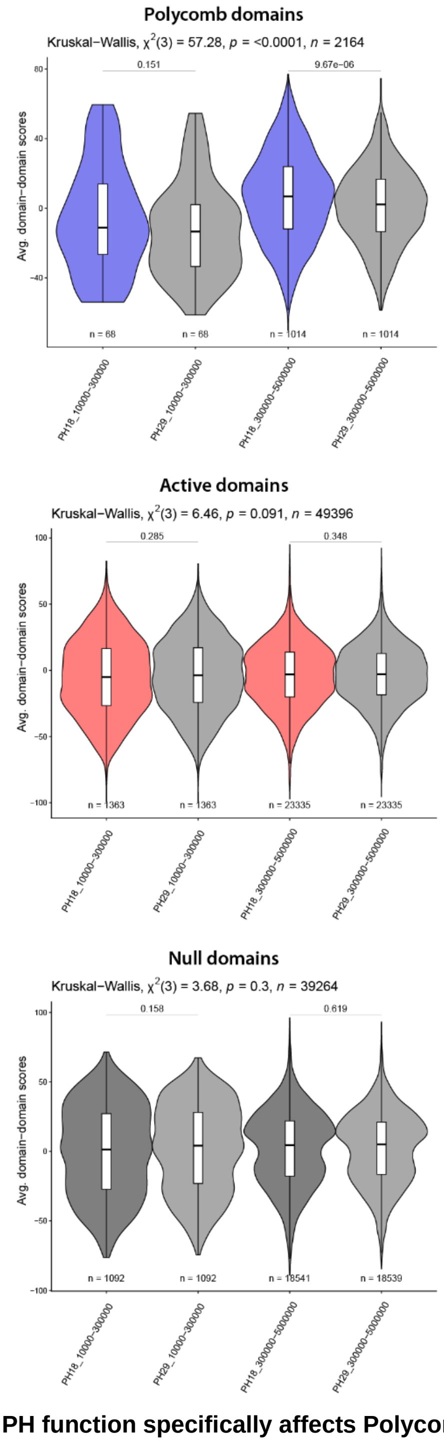


**Supplementary Figure 6: Loss of PH function specifically affects Polycomb domain interactions.** Violin plot shows the quantification of the average score between PcG (*top*), Active (*middle*), and Null (*bottom*) domains identified using ChromHMM (https://ernstlab.github.io/ChromHMM/) (45) at 20kb resolution from ATAC-seq, H2AK118Ub, H3K27ac, and H3K27me3 signals in eye disc Control samples. Inter-domain scores are classified in shorter (from 10 to 300 kb) and longer-range (from 300 kb to 5 Mb) of genomic separation. Statistical comparisons were performed using the Kruskal–Wallis test followed by Dunn's post-hoc pairwise test with Benjamini–Hochberg correction. Boxplots show median (central line), Q1=25th and Q3=75th percentiles (box limits), and Q1+1.5×IQR to Q3+1.5×IQR (whiskers), where IQR is the interquartile range.

#### Figure S7

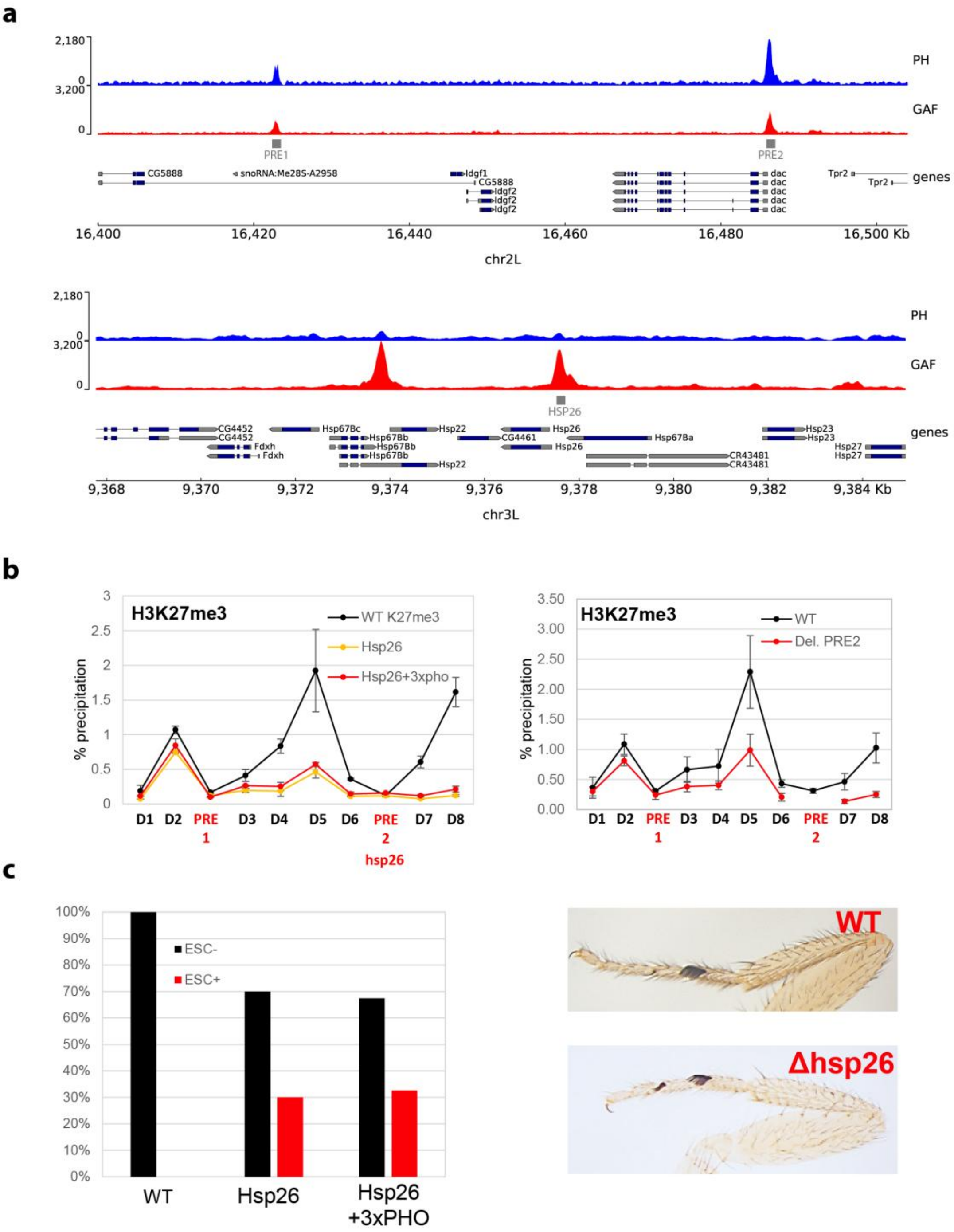


**Supplementary Figure 7: Hsp26 sequences do not recruit PcG proteins in the context of a PcG domain**
**(a)** CUT&RUN PH and GAF profiles at the *dac* gene locus on chromosome 2L (top), or the *hsp26* gene locus on chromosome 3L (bottom). **(b)** qChIP experiments using H3K27me3 antibody in WT, Hsp26, and Hsp26+3xPHO fly lines (left), or ΔPRE2 line (right). D1-D8, and PRE1, 2 indicate PCR amplicons used for qChIP experiments along the *dac* gene locus (**Supplementary Table 2**). The housekeeping gene *Rp49* was used as negative control, the *engrailed* PRE as a positive control. Data are presented as the mean values ± s.d (error bars) of two independent replicates. **(c)** Quantification of the extra sex comb (ESC) phenotype in the indicated fly lines grown at 25°C. A minimum of 50 male flies were scored (left). Representative examples of a WT and mutant phenotype leg are shown (right).

#### Figure S8

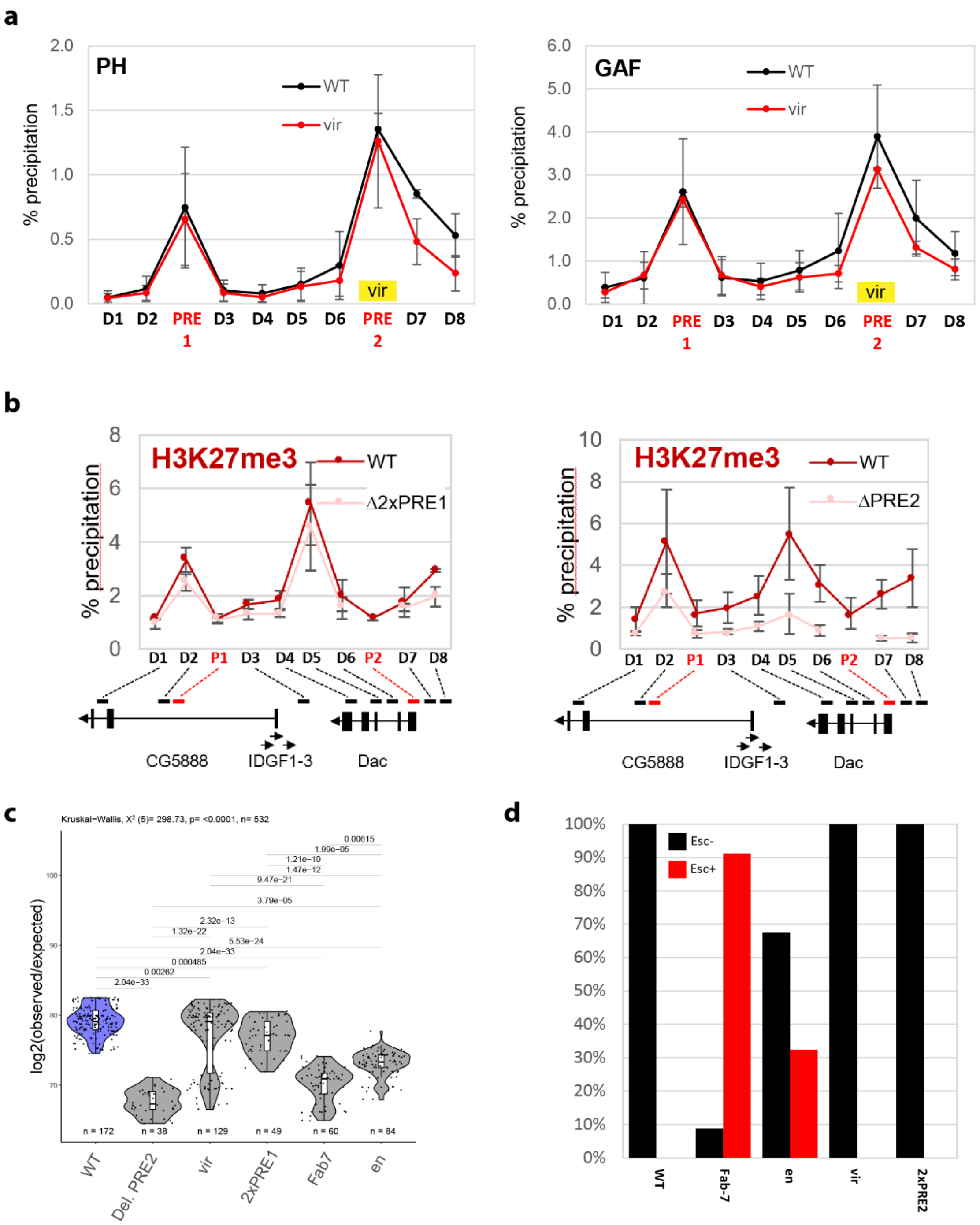


**Supplementary Figure 8: The orthologous PRE sequence from *D. virilis* recruits GAF and PH proteins in *D. melanogaster***

**(a)** qChIP experiments using PH or GAF antibodies in WT flies or PRE-mutant flies where *dac* PRE2 has been replaced by the orthologous sequence from *D. virilis* (vir). **(b)** qChIP experiments using H3K27me3 antibody in WT flies or PRE-mutant flies where *dac* PRE2 has been replaced by PRE1 (2xPRE1) or deleted (ΔPRE2). D1-D8, and PRE1, 2 indicate PCR amplicons used for qChIP experiments along the *dac* gene locus in panels **(a)** and **(b)** (**Supplementary Table S2**). The housekeeping gene *Rp49* was used as negative control, the *engrailed* PRE as a positive control. Data are presented as the mean values ± s.d (error bars) of two independent replicates. **(c)** Hi-C interaction score at the *dac* PRE in WT, ΔPRE2, and mutant PRE lines, where the *dac* PRE2 has been replaced by the indicated PREs (vir or 2xPRE1 and Fab-7, en). The number of points per distribution is reported (**Material and Methods**). Boxplots show median (central line), Q1=25th and Q3=75th percentiles (box limits), and Q1+1.5×IQR to Q3+1.5×IQR (whiskers), where IQR is the interquartile range. Statistical comparisons were performed using the Kruskal–Wallis test followed by Dunn's post-hoc pairwise test with Benjamini–Hochberg correction. **(d)** Quantification of the extra sex comb (ESC) phenotype in the indicated fly lines grown at 25°C. A minimum of 50 male flies were scored.

**Supplementary Table 1:** Sequencing statistics and processed data of the CUT&RUN, Micro-C and Hi-C samples generated for this work.

**Supplementary Table 2:** Primer sequences used for the generation of fly lines and qCHIP
